# Interdependence of cellular and network properties in respiratory rhythmogenesis

**DOI:** 10.1101/2023.10.30.564834

**Authors:** Ryan S. Phillips, Nathan A. Baertsch

## Abstract

How breathing is generated by the preBötzinger Complex (preBötC) remains divided between two ideological frameworks, and the persistent sodium current (*I*_*NaP*_) lies at the heart of this debate. Although *I*_*NaP*_ is widely expressed, the *pacemaker hypothesis* considers it essential because it endows a small subset of neurons with intrinsic bursting or “pacemaker” activity. In contrast, *burstlet theory* considers *I*_*NaP*_ dispensable because rhythm emerges from “pre-inspiratory” spiking activity driven by feed-forward network interactions. Using computational modeling, we discover that changes in spike shape can dissociate *I*_*NaP*_ from intrinsic bursting. Consistent with many experimental benchmarks, conditional effects on spike shape during simulated changes in oxygenation, development, extracellular potassium, and temperature alter the prevalence of intrinsic bursting and pre-inspiratory spiking without altering the role of *I*_*NaP*_. Our results support a unifying hypothesis where *I*_*NaP*_ and excitatory network interactions, but not intrinsic bursting or pre-inspiratory spiking, are critical interdependent features of preBötC rhythmogenesis.

**SIGNIFICANCE STATEMENT:** Breathing is a vital rhythmic process originating from the preBötzinger complex. Since its discovery in 1991, there has been a spirited debate about whether respiratory rhythm generation emerges as a network property or is driven by a subset of specialized neurons with rhythmic bursting capabilities, endowed by intrinsic currents. Here, using computational modeling, we propose a unifying data-driven model of respiratory rhythm generation which bridges the gap between these competing theories. In this model, both intrinsic cellular properties (a persistent sodium current) and network properties (recurrent excitation), but not intrinsic bursting, are essential and interdependent features of respiratory rhythm generation.

## INTRODUCTION

Neural rhythmicity orchestrates critical brain functions (Wang, 2010; Fries, 2023; Başar and Düzgün, 2016; Guan et al., 2022) and dysregulation of this rhythmicity can lead to pathology (Stafstrom, 2007; Hammond et al., 2007). Due to their experimental accessibility, central pattern generators (CPGs) that drive vital invertebrate and vertebrate rhythmic functions such as locomotion and digestion have served as key model systems for investigating how the brain generates rhythm (Selverston, 2010; Daur et al., 2016; MacKay-Lyons, 2002; Marder and Bucher, 2001; Marder et al., 2005). In mammals, the CPG for breathing has been perhaps the most extensively studied as this network produces a vital motor output that can be readily measured in awake, anesthetized, and *ex vivo* experimental preparations. Discovery the preBötzinger Complex (preBötC), a region in the ventrolateral medulla that is necessary for respiratory rhythm, inspired the development of slice preparations from neonatal rodents (Smith et al., 1991; Ramirez et al., 1996; Johnson et al., 2001; Funk and Greer, 2013) that capture enough of this network for it to continue to generate rhythm when isolated from the rest of the brain. These *in vitro* preparations have been used extensively over the last three decades in an ongoing effort to identify properties of the preBötC that underlie rhythmogenesis. Computational modeling studies conducted in parallel have been critical for testing concepts that are experimentally intractable and for developing new predictions for subsequent experimental (in)validation. Yet, despite rigorous experimental/theoretical investigation and the deceptive simplicity of breathing, how the preBötC network generates rhythm remains controversial and unresolved (Feldman and Del Negro, 2006; Feldman and Kam, 2015; Molkov et al., 2017; Ramirez and Baertsch, 2018; Ashhad et al., 2022; Smith, 2022).

The terminology surrounding this controversy has evolved since first being formally discussed (Rekling and Feldman, 1998). However, the overall nature of the debate has remained centered on whether cellular- or network-based properties of the preBötC are the essential mechanism of rhythm generation. Much of the contemporary debate relates to two competing theories. With its discovery, preBötC neurons were identified that continue to produce rhythmic bursts of action potentials following pharmacological blockade of synaptic interactions (Smith et al., 1991). This finding, as well as observations that attenuation of synaptic inhibition does not block the respiratory rhythm (Rekling and Feldman, 1998; Johnson et al., 2001), inspired the *pacemaker hypothesis*, which posits that these intrinsically bursting neurons or “pacemakers” are a specialized group of neurons that initiates synchronized activity within the network and represent the essential element of rhythmogenesis. Computational modeling studies predicted a critical role of a slowly inactivating persistent sodium current (*I*_*NaP*_) in the intrinsic oscillatory activity of pacemaker neurons (Butera et al., 1999a), which was later experimentally confirmed (Del Negro et al., 2002a; Koizumi and Smith, 2008; Yamanishi et al., 2018). More recently, an alternative view has evolved to account for observations that the amplitude of the preBötC rhythm can be diminished while only minimally affecting its frequency (Johnson et al., 2001; Del Negro et al., 2002b; Peña et al., 2004; Pace et al., 2007a; Koizumi et al., 2018; Picardo et al., 2019; Del Negro et al., 2001, 2009; Koizumi et al., 2016; Sun et al., 2019; Phillips et al., 2022), suggesting that the network contains dissociable rhythm and “burst” generating elements (Kam et al., 2013b; Phillips et al., 2019, 2022; Ashhad and Feldman, 2020; Phillips and Rubin, 2022). One interpretation of these results is conceptualized as *burstlet theory* (Feldman and Kam, 2015), based on elements of the preceding “*group pacemaker*” hypothesis (Rekling and Feldman, 1998), which proposes that rhythm is driven by weakly synchronized spiking activity referred to as “burstlets” that are an emergent property of preBötC network topology and feed-forward excitatory synaptic interactions among a subset of non-pacemaker neurons. Thus, in burstlet theory, ramping spiking activity prior to the onset of inspiratory bursts referred to as “pre-inspiratory spiking” represents the burstlet and is the essential rhythmogenic element of the network, while intrinsic bursting neurons and associated burst-promoting conductances including *I*_*NaP*_ are considered dispensable (Del Negro et al., 2002b, 2005; Feldman and Del Negro, 2006; Feldman and Kam, 2015; Ashhad et al., 2022; da Silva et al., 2019).

However, both theories are difficult to test using experimental approaches and are limited by conflicting findings and oversimplifications that have hindered progress toward consensus on how breathing originates. Initially, the *pacemaker hypothesis* was widely adopted due to its simplicity and convincing experimental (Smith et al., 1991; Johnson et al., 1994; Koshiya and Smith, 1999; Del Negro et al., 2002a; Koizumi and Smith, 2008) and theoretical (Butera et al., 1999a,b; Del Negro et al., 2001) support. Yet, despite its appeal, demonstrating that intrinsic bursting neurons are critical for rhythmogenesis proved to be far from simple. First, intrinsic bursting neurons are difficult to identify, typically requiring blockade of synaptic network interactions rendering the network non-functional. Second, even if identifiable in the active network, intrinsic bursting neurons cannot be specifically manipulated to define their functional role. For example, although *I*_*NaP*_ is higher on average in intrinsic bursters, *I*_*NaP*_ is widely expressed in the preBötC in both intrinsically bursting and non-bursting neurons (Del Negro et al., 2002a; Ptak et al., 2005; Koizumi and Smith, 2008; Yamanishi et al., 2018). Because of this ubiquitous expression, manipulations of *I*_*NaP*_ are not specific to intrinsic bursting neurons making it difficult or impossible to characterize their specific contribution to rhythm generation. Third, intrinsic bursting does not appear to be a fixed property of the preBötC network since neurons can be capable of bursting in some conditions, but not in others (Hilaire and Duron, 1999; Smith et al., 2000; Massey et al., 2014; Ptak et al., 2009; Peña and Ramirez, 2002). For instance, when challenged with hypoxia, the preBötC network produces a gasping-like rhythm that has enhanced sensitivity to *I*_*NaP*_ blockade (Peña et al., 2004; Paton et al., 2006) and is associated with a loss of pre-inspiratory spiking, inconsistent with the rhythmogenic mechanism proposed by *burstlet theory*. On the other hand, identification of preBötC pacemaker neurons has relied on *ex vivo* preparations from neonatal mice with associated caveats such as elevated extracellular [*K*^+^] and low temperature (Smith et al., 1991; Johnson et al., 1994; Del Negro et al., 2001; Koizumi and Smith, 2008; Yamanishi et al., 2018; Phillips et al., 2022), while there remains a lack of evidence for intrinsically bursting neurons in adult animals *in vivo*, casting doubt on the *pacemaker hypothesis* (Feldman and Del Negro, 2006; Feldman and Kam, 2015; Ashhad et al., 2022).

Here, we develop a new model of respiratory rhythmogenesis that accounts for these discrepancies, while remaining constrained by experimental findings that support both the *pacemaker hypothesis* and *burstlet theory*. Due to interactions with the voltage-dependent properties of *I*_*NaP*_, we find that small changes in spike shape, without changes in *I*_*NaP*_ expression or excitability, can eliminate the capability of model neurons to exhibit intrinsic bursting. By exploiting this interaction to dissociate the role of *I*_*NaP*_ from the role of intrinsic bursting in model preBötC networks, we find that networks comprised entirely of neurons that are rendered incapable of intrinsic bursting continue to produce rhythm. In this extreme case, excitatory synaptic interactions allow rhythm to emerge among tonic neurons that typically exhibit pre-inspiratory spiking in the synaptically coupled network. Yet, despite the absence of intrinsic bursting, rhythm generation in these networks remains dependent on *I*_*NaP*_. At the other extreme, in networks with spike shapes that render all neurons capable of intrinsic bursting, rhythmogenesis continues despite minimal pre-inspiratory spiking. In this case, the network rhythm also depends on *I*_*NaP*_ as well as excitatory interactions that synchronize intrinsic bursting to produce a coherent network rhythm. Introducing spike shape variability allows subsets of neurons to regain intrinsic bursting capabilities or pre-inspiratory spiking, but this does not endow them with a specialized role in rhythm generation *per se*. Instead, the interdependence of *I*_*NaP*_ and excitatory synaptic interactions represents the critical substrate for rhythmogenesis, while intrinsic bursting and pre-inspiratory spiking are conditional phenotypes of preBötC neurons sensitive to any perturbation that affects spike shape, including, but not limited to, extracellular [K+], temperature, hypoxia, and neurodevelopment. These findings support a unifying theory of respiratory rhythm generation and may also provide a useful framework for understanding the emergence of rhythmicity in other brain networks.

## RESULTS

### Spike shape regulates intrinsic bursting

Spike shapes vary widely, even within specific brain regions (Bean, 2007). In the preBötC, spike heights can range from approximately 15 − 20 *mV* (Thoby-Brisson and Ramirez, 2001; Tryba et al., 2003; Peña and Ramirez, 2004; Zavala-Tecuapetla et al., 2008) to 100 − 125 *mV* (Koshiya and Smith, 1999; Del Negro et al., 2001; Krey et al., 2010). Due to the voltage-dependence of *I*_*NaP*_ (in)activation (Del Negro et al., 2002a; Ptak et al., 2005; Koizumi and Smith, 2008; Yamanishi et al., 2018), we wondered whether spike shape could impact intrinsic bursting. To selectively manipulate spike shape, we incorporated two additional currents, *I*_*SPK*_ and *I*_*AHP*_, into a contemporary preBötC neuron model (Phillips and Rubin, 2022), (Fig. 1A). The voltage-dependent properties of *I*_*SPK*_ and *I*_*AHP*_ (in)activation were chosen such that they are only active well above resting membrane potential, allowing selective control of spike shape without affecting excitability. Although not intended to mimic any one of the numerous ion channels expressed in the preBötC that may influence spike shape (Ptak et al., 2005; Krey et al., 2010; Phillips et al., 2018; Revill et al., 2021; Burgraff et al., 2022), the voltage-dependent properties of *I*_*SPK*_ and *I*_*AHP*_ are similar to NaV1.2 (Plant et al., 2016) and non-inactivating M-currents (Manville and Abbott, 2019), respectively, that are expressed by preBötC neurons (Ptak et al., 2005; Revill et al., 2021). For a full model description see *Materials and Methods*. As expected, increasing the *I*_*SPK*_ conductance (*g*_*SPK*_) increased spike height by ≈ 30 *mV* (− 18.12 *mV* to +11.22 *mV*) over the range of conductances tested (0 − 50 *nS*). Over the same range of *g*_*SPK*_, the magnitude of the spike afterhyperpolarization (AHP) was also increased by ≈ 6 *mV* (− 55.14 *mV* to − 61.13 *mV*). In contrast, increasing the *I*_*AHP*_ conductance (*g*_*AHP*_) from 0 *nS* to 50 *nS* had a more selective effect on spike AHP, increasing it by a similar amount from − 55.14 *mV* to − 60.63 *mV* with minimal changes in spike height (Fig. 1B & C).

**Figure 1.**
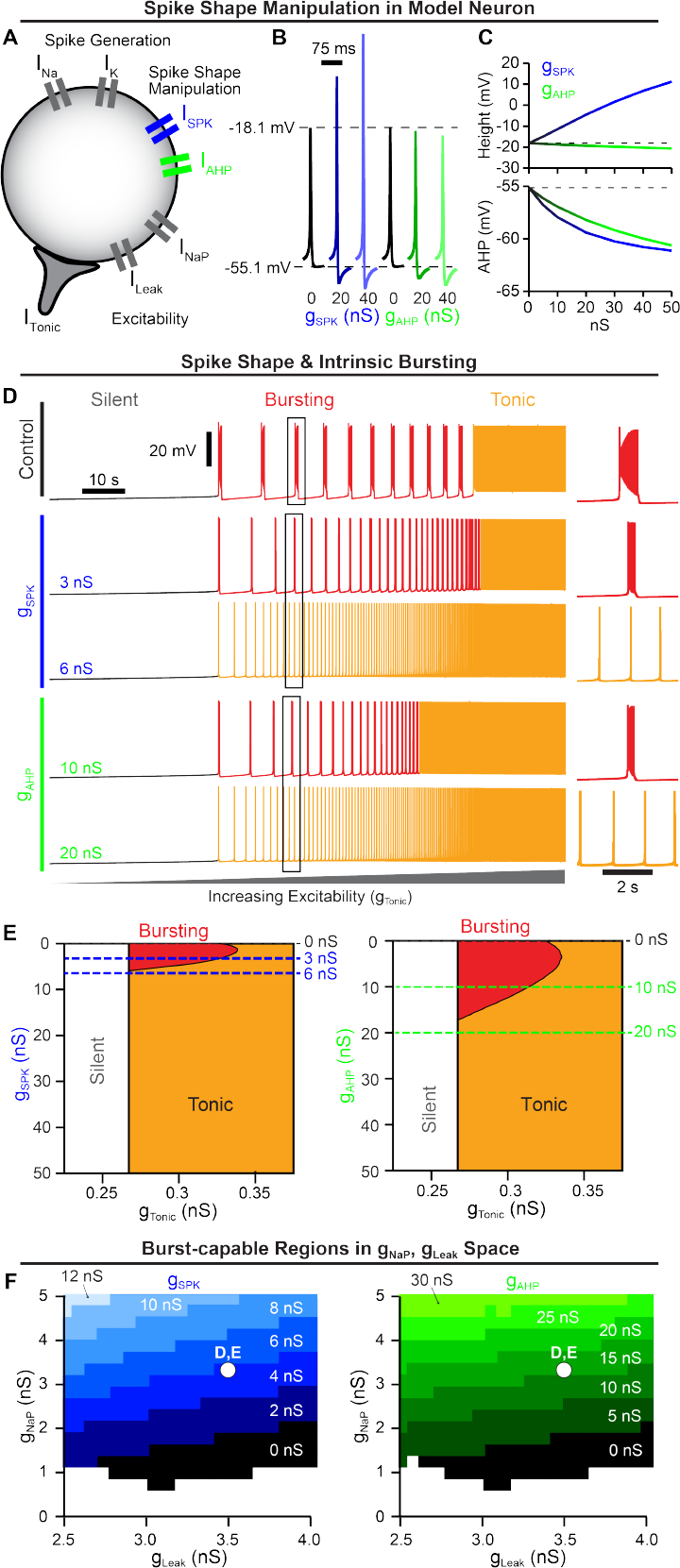
Spike shape regulates *I*_*NaP*_-dependent intrinsic bursting. (A) Schematic diagram of a model neuron with modifiable spike shape. (B) Example spike shapes and (C) quantification of spike height and AHP during increasing *g*_*SPK*_ or *g*_*AHP*_. (D) Voltage traces of an average intrinsic burster (*g*_*NaP*_ = 3.33 *nS* and *g*_*leak*_ = 3.5 *nS* (Koizumi and Smith, 2008)) illustrating how increasing *g*_*SPK*_ or *g*_*AHP*_ changes the activity pattern (silent, bursting, or tonic) produced as *g*_*Tonic*_ is varied. (E) Activity patterns as a function of *g*_*Tonic*_ and *g*_*SPK*_ (left) or *g*_*AHP*_ (right). Notice that for small increases in *g*_*SPK*_ or *g*_*AHP*_, intrinsic bursting (red shaded region) is lost and the neuron is rendered ‘burst-incapable’. (F) burst-capable regions of *g*_*Leak*_, *g*_*NaP*_ space as *g*_*SPK*_ (left) or *g*_*AHP*_ (right) is increased. White dot indicates *g*_*Leak*_, *g*_*NaP*_ values of neuron in D & E.

To characterize the interaction between spike shape and intrinsic bursting, we altered *g*_*SPK*_ or *g*_*AHP*_ in model neurons with experimentally motivated *I*_*NaP*_ conductance (*g*_*NaP*_) (Del Negro et al., 2002a; Koizumi and Smith, 2008; Koizumi et al., 2010; Yamanishi et al., 2018) while manipulating excitability via a tonic excitatory conductance *g*_*Tonic*_. Importantly, intrinsic bursting is voltage-dependent as illustrated in Fig. 1D (top) where the model neuron transitions from silent, to intrinsic bursting (periodic bursts of spiking), and then to tonic spiking (continuous spiking) as excitability increases (Smith et al., 1991; Butera et al., 1999a; Del Negro et al., 2002a; Koizumi and Smith, 2008). Surprisingly, we found that small increases in spike height or AHP rapidly reduced the range of excitability (*g*_*Tonic*_) where intrinsic bursting was possible, followed by complete elimination of intrinsic bursting capabilities at *g*_*SPK*_ = 5.816 *nS* or *g*_*AHP*_ = 17.143 *nS* corresponding to changes in spike height or AHP of approximately +3.5 *mV* and − 3.0 *mV*, respectively (Fig. 1D & E). Specifically, as spike height or AHP increased with excitability held constant, the duration and period of intrinsic bursts were reduced as the number of spikes and their frequency during bursts decreased until the neuron transitioned to tonic spiking (Fig. 1D right insets and Fig. 1-Supplement 1). Importantly, following these changes in spike shape, neurons remained unable to generate intrinsic bursting at all levels of excitability, transitioning directly from silent to tonic spiking, which we refer to here as being “burst-incapable”. Notably, because this designation as burst-capable or -incapable accounts for all levels of excitability, it is distinct from the more common terminology referring to the voltage-dependent transition in or out of an intrinsic bursting “mode”. In addition to *g*_*NaP*_, the potassium-dominated leak conductance (*g*_*Leak*_) is an important determinant of intrinsic bursting properties and varies among preBötC neurons (Del Negro et al., 2002a; Koizumi and Smith, 2008; Yamanishi et al., 2018). Therefore, we mapped burst capability across *g*_*NaP*_,*g*_*Leak*_ parameter space during manipulations of spike shape (Fig. 1F). As *g*_*SPK*_ or *g*_*AHP*_ were increased, the burst-capable region collapsed towards higher *g*_*NaP*_ and lower *g*_*Leak*_ values until intrinsic bursting became impossible at values of *g*_*SPK*_ or *g*_*AHP*_ greater than 13 *nS* or 35 *nS*, respectively. Thus, even model neurons with high *g*_*NaP*_ and low *g*_*Leak*_ require spike shape to be maintained within a certain range to be capable of intrinsic bursting.

### Intrinsic bursting is not required for preBötC network rhythmogenesis

Demonstrating a critical role of intrinsic bursting or “pacemaker” neurons for rhythm generation in the preBötC and other CPGs has been difficult (see *Introduction*) and controversial (Smith et al., 2000; Feldman and Kam, 2015; Molkov et al., 2017; Del Negro et al., 2018; Ashhad et al., 2022; Smith, 2022). Therefore, we leveraged the interaction between spike shape and intrinsic bursting described above to investigate how manipulation of intrinsic bursting, without associated changes in *I*_*NaP*_ or excitability, impacts rhythm generation in a network of *N* = 100 model neurons (Fig. 2A). Because the preBötC contains rhythm- and pattern (burst amplitude)-generating subpopulations (Kam et al., 2013a; Phillips et al., 2019; Sun et al., 2019; Phillips and Rubin, 2022; Phillips et al., 2022; Ashhad and Feldman, 2020), our model network is intended to represent the rhythm-generating subpopulation (≈ 25% of preBötC neurons) thought to be enriched in intrinsic bursters and neurons with pre-inspiratory spiking activity (Rekling and Feldman, 1998). The parameters of the model network are data-driven, using experimentally motivated synaptic connectivity probability (13%) (Rekling et al., 2000), synaptic depression (Kottick and Del Negro, 2015), and distributions of *g*_*NaP*_ and *g*_*Leak*_ (Del Negro et al., 2002a; Koizumi and Smith, 2008), as initially described in (Phillips and Rubin, 2022) (Fig. 2A & B). See *Material and Methods* for a full model description.

**Figure 2.**
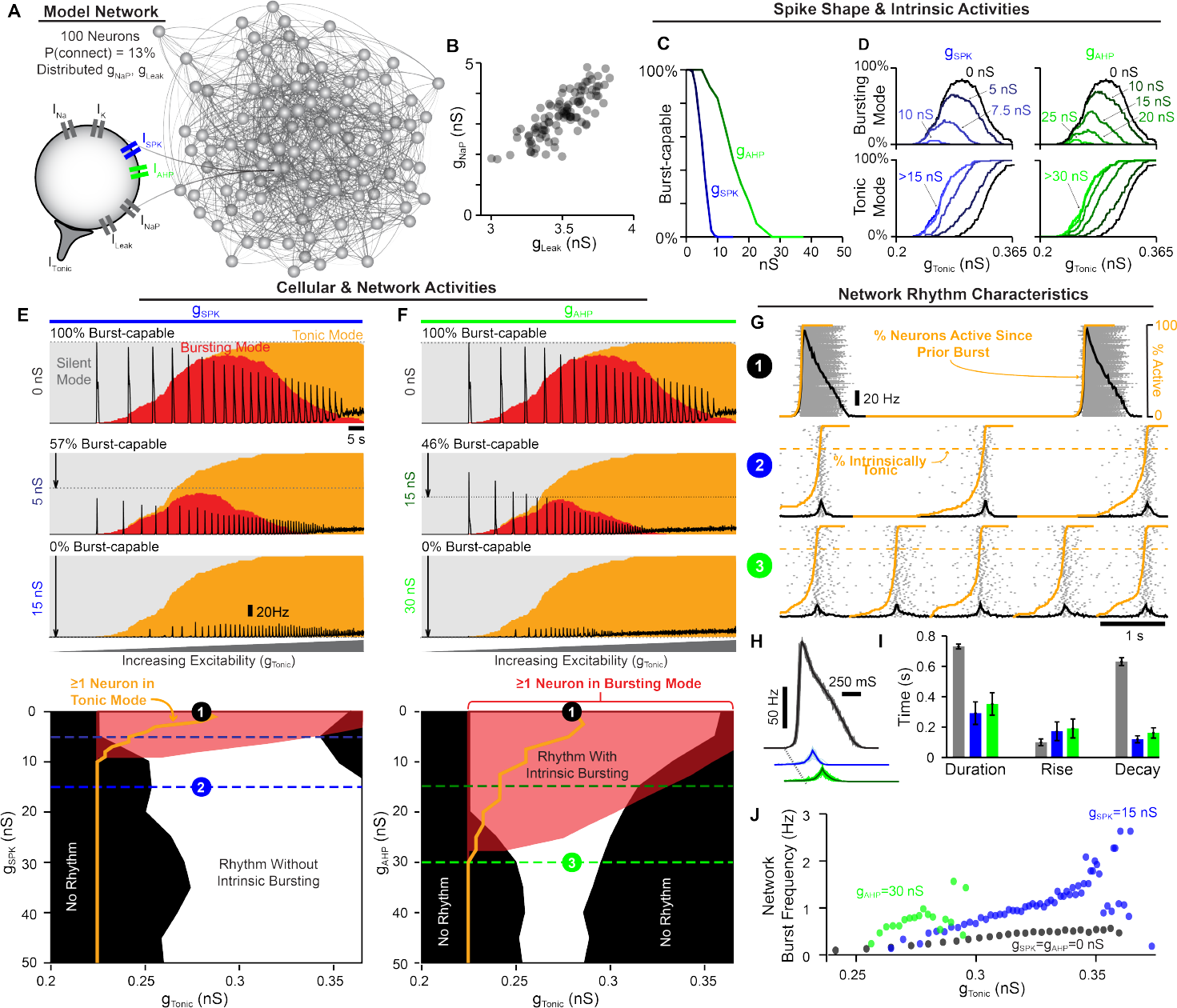
Rhythm generation continues following spike-shape-induced elimination of intrinsic bursting. (A) Schematic of 100 neuron network. (B) Distribution of *g*_*NaP*_ and *g*_*Leak*_ within the example network. (C) Percentage of the network that is burst-capable as a function of *g*_*SPK*_ or *g*_*AHP*_. (D) Relationship between *g*_*Tonic*_ and the percentage of the population in bursting (top) or tonic (bottom) modes during increasing *g*_*SPK*_ (left) or *g*_*AHP*_ (right). Effects of increasing (E) *g*_*SPK*_ or (F) *g*_*AHP*_ on the network activity (firing rate) and intrinsic cellular activity modes (silent, bursting, tonic) as excitability is increased with corresponding parameter space supporting intrinsic bursting (red), tonic spiking (orange lines), and network rhythmogenesis (white) shown below. Dotted lines correspond to example traces. (G) Example raster plots with overlaid population firing rate for each condition at a fixed *g*_*Tonic*_. (H) Cycle-triggered averages of network burst waveforms and (I) quantification of burst duration, rise, and decay times. (J) Effect of increasing *g*_*SPK*_ or *g*_*AHP*_ on the range of possible network burst frequencies.

Because these model networks contain neurons with distributed *g*_*NaP*_ and *g*_*Leak*_ values, we first characterized how increasing spike height and/or spike AHP impacts intrinsic bursting capabilities across the population. Under control spike shape conditions (*g*_*SPK*_ = *g*_*AHP*_ = 0 *nS*), all neurons were initially burst-capable. However, due to the spike shape dependence of bursting capabilities (see Fig. 1, increasing *g*_*SPK*_ or *g*_*AHP*_ progressively rendered neurons incapable of intrinsic bursting, with low *g*_*NaP*_ neurons being the most susceptible. When spike height was increased by as little as ≈ 10 *mV* (*g*_*SPK*_ = 10 *nS*) or the AHP was increased by ≈ 4 *mV* (*g*_*AHP*_ = 30 *nS*), the intrinsic bursting capabilities of all neurons in the population were eliminated, i.e. they became burst-incapable. (Fig. 2C).

Next, we examined the activity of the synaptically coupled network in relation to the intrinsic activity modes (silent, bursting, tonic) of its constituent neurons. This was done by determining the percentage of neurons in the network that are silent, bursting, or tonic in the absence of synaptic interactions as a function of excitability (*g*_*Tonic*_) (Fig. 2D). Excitatory synaptic interactions were then introduced, and population firing rate over the same excitability range was overlaid with intrinsic activity modes to compare cellular- and network-level characteristics (Fig. 2E & F). Under control spike shape conditions (100% burst-capable), as excitability was increased the percentage of neurons in an intrinsic bursting mode increased and then decreased as neurons transitioned to tonic mode. This revealed a bell-shaped curve where the maximum number of neurons in bursting mode was always less than the number of burst-capable neurons. This occurs because the *g*_*NaP*_, *g*_*Leak*_ parameters of each neuron are drawn from a distribution, and therefore not all burst-capable neurons are in bursting mode at a given level of *g*_*Tonic*_. Following introduction of synaptic connections, the control network produced a rhythm that followed this bell-shaped curve, beginning as soon as the first neuron entered an intrinsic bursting mode and ending once most neurons switched to tonic mode. In model networks with altered spike shape, the bell-shaped curve of neurons in bursting mode was initially reduced, involving a smaller percentage of the network and occurring over a narrower range of excitability, and then eliminated once all neurons were rendered burst-incapable at *g*_*SPK*_ ≈ 10 *nS* or *g*_*AHP*_ ≈ 30 *nS*, as described above. Consequently, as excitability was increased, neurons gradually transitioned directly from silent to tonic modes. Surprisingly, with synaptic connections introduced, the network still became rhythmic once a sufficient fraction of neurons (≈ 15%) entered tonic mode. Thus, contrary to the *pacemaker hypothesis* and the expected mechanism of rhythm generation in similar *I*_*NaP*_-based model networks (Butera et al., 1999b; Del Negro et al., 2001; Jasinski et al., 2013; Phillips et al., 2019, 2022; Phillips and Rubin, 2022), intrinsic bursting is not required for rhythm generation.

Next, we examined how changes in spike shape impact patterns of population spiking activity, Fig. 2G. Altering either spike shape feature reduced the amplitude of the network rhythm due to a decrease in the firing rates of individual neurons during bursts to ≈ 20 *Hz*. These network rhythms may appear relatively weak; however, the much larger amplitude rhythm under control conditions, with spike rates reaching *>* 130 *Hz* in many neurons, is less representative of preBötC activity since spike rates of preBötC neurons during bursts typically range from very slow (*<* 5*Hz*) to a maximum near ≈ 50*Hz* (Kam et al., 2013a; Johnson et al., 1994; Yamanishi et al., 2018; Baertsch and Ramirez, 2019). Increasing *g*_*SPK*_ or *g*_*AHP*_ also converted network bursts from a decrementing pattern to one with roughly symmetrical rise and decay times on the order of 150− 200 *ms* (Fig. 2H & I), which is also more representative of typical preBötC activity and compatible with the cellular and network level dynamics of burstlet oscillations (Kallurkar et al., 2020; Kam et al., 2013a). In addition to altered firing patterns during bursts, modifying spike shape led to changes in the spiking activity of neurons between bursts. Specifically, at a given level of *g*_*Tonic*_, the fraction of neurons in the network that began to spike prior to burst onset became much larger when *g*_*SPK*_ or *g*_*AHP*_ was increased, resulting in a collective “pre-inspiratory” ramping of network activity (orange lines in Fig. 2G). As suggested by previous recordings of preBötC neurons (Butera et al., 1999b; Kam et al., 2013a; Baertsch et al., 2021; Kallurkar et al., 2020), this pre-inspiratory activity in the model network reflects the recovery of activity in neurons that are in tonic spiking mode Fig. 2-Supplement 2.

Altering spike shape to reduce or eliminate intrinsic bursting also changed how the network rhythm responded to modulation of excitability. Specifically, the *g*_*Tonic*_ range supporting rhythmogenesis was altered slightly with increasing *g*_*SPK*_ and reduced with *g*_*AHP*_, (Fig. 2E & J). Yet, despite the reduced excitability “window”, the responsiveness of the network to changes in *g*_*Tonic*_ was enhanced such that the dynamic range of possible burst frequencies increased by 2-3 fold. Further, in networks lacking intrinsic bursting, the window of excitability sufficient to produce rhythm could be substantially increased by increasing synaptic strength (Fig. 2-Supplement 1). Overall, these results demonstrate that 1) rhythmogenesis can persist even in the extreme case when all neurons are rendered incapable of intrinsic bursting, 2) reducing the number of burst-capable neurons without altering *I*_*NaP*_ produces a network rhythm with spiking patterns that are more representative of preBötC activity, and 3) modulation of spike height can change the gain of the network rhythm such that it responds with a greater change in frequency to a given excitatory input.

### Interdependence of *I*_*NaP*_ and excitatory synaptic dynamics

Our finding that rhythmogenesis continues without intrinsic bursting was surprising since *I*_*NaP*_-based computational models of the preBötC are generally viewed as the embodiment of the *pacemaker hypothesis*. In other preBötC models that lack *I*_*NaP*_ (and intrinsic bursting as a result), specialized synaptic dynamics (depression/facilitation) can underlie network oscillations (Rubin et al., 2009; Guerrier et al., 2015). Similarly, synapses in our preBötC model undergo activity-dependent synaptic depression as motivated by experimental observations (Kottick and Del Negro, 2015). Therefore, to better understand what underlies rhythm generation in the model network, we blocked *I*_*NaP*_ or removed synaptic depression under control conditions with 100% burst-capable neurons (*g*_*SPK*_ = *g*_*AHP*_ = 0 *nS*) and also following elimination of intrinsic bursting via increased spike height and/or AHP (*g*_*SPK*_ = 15 *nS* or *g*_*AHP*_ = 35 *nS*). Under all conditions, network rhythms continued when synaptic depression was turned off, with modestly increased burst duration and decreased burst frequency (Fig. 3A). Surprisingly, in the absence of synaptic depression, the excitability range supporting rhythmogenesis was substantially reduced in control networks with 100% burst-capable neurons but slightly increased in networks lacking intrinsic bursting. Thus, in the model network, synaptic depression has important effects on rhythm characteristics, but its elimination does not preclude rhythmogenesis.

**Figure 3.**
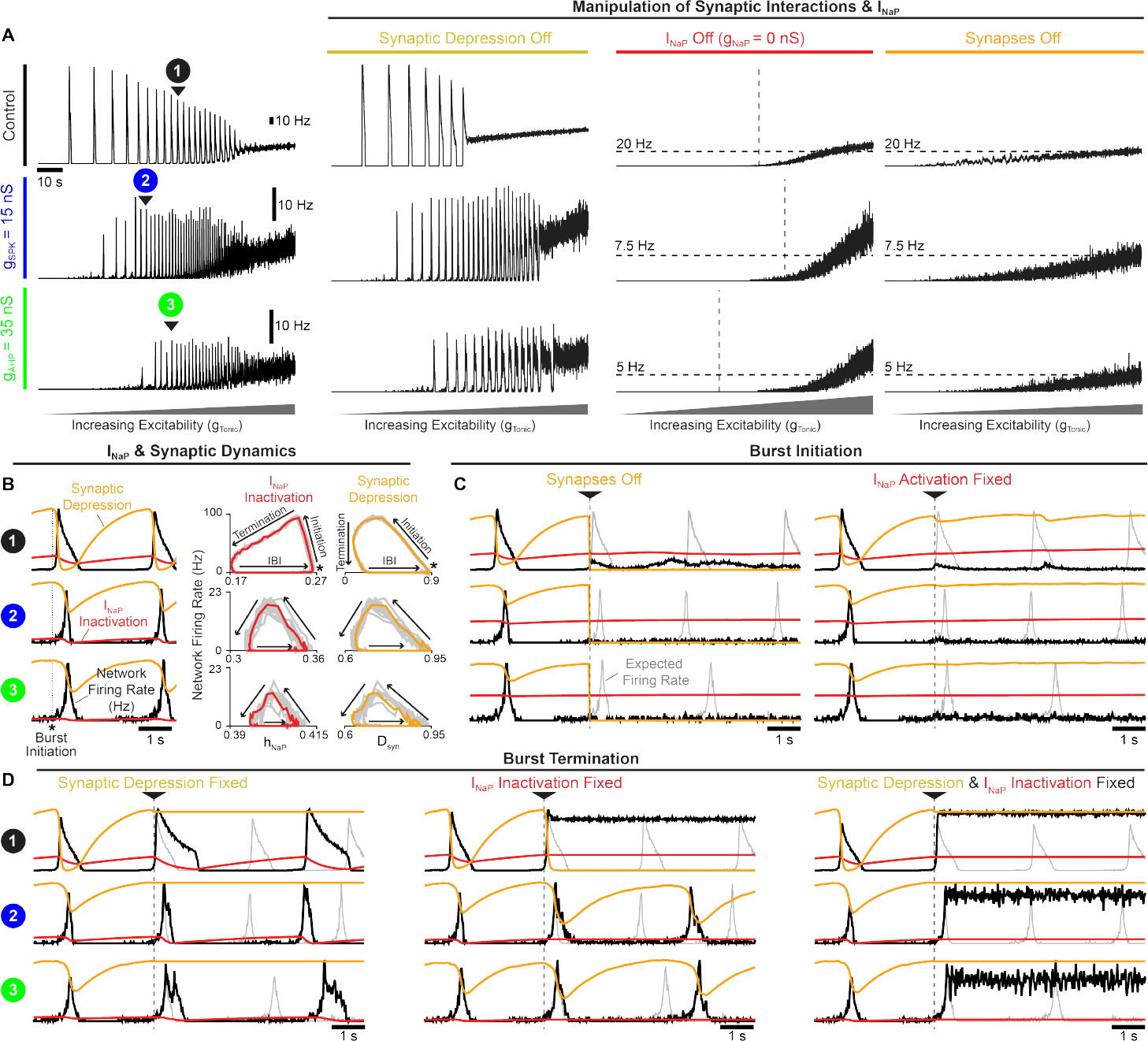
Interdependence of *I*_*NaP*_ and synaptic interactions for network rhythmogenesis. (A) Activity of networks with all burst-capable (control) or burst-incapable (*g*_*SPK*_ = 15 *nS* or *g*_*AHP*_ = 35 *nS*) neurons following elimination of synaptic depression, *I*_*NaP*_, or all synaptic interactions. (B) Relationship between network firing rate, *I*_*NaP*_ inactivation, and synaptic depression during network burst initiation, termination, and the inter-burst interval. (C) Network activity when synapses are turned off (left) or *I*_*NaP*_ activation (*m*_*NaP*_) is fixed in neurons that have not yet spiked (right) at burst initiation. (D) Network activity when synaptic depression (left), *I*_*NaP*_ inactivation (*h*_*NaP*_, middle), or both (right), are fixed at burst initiation. Gray traces indicate expected network activity, orange traces represent synaptic depression, and red traces indicate *I*_*NaP*_ inactivation.

To explore the role of *I*_*NaP*_, we set *g*_*NaP*_ = 0 *nS* to eliminate its activity from all neurons in the network. As expected, under all conditions (control, *g*_*SPK*_ = 15 *nS, g*_*AHP*_ = 35 *nS*), removing *I*_*NaP*_ decreased neuronal excitability resulting in higher levels of *g*_*Tonic*_ required to drive spiking activity. However, all networks remained unable to produce rhythm even as *g*_*Tonic*_ was increased to restore excitability to levels that produced comparable spike rates (Fig. 3A). For comparison, networks with synapses blocked (*g*_*syn*_ = 0 *ns*) were also unable to produce rhythm at any level of excitability, illustrating the somewhat trivial but important point that synaptic interactions are always a requirement for network rhythm, even if all neurons are intrinsic bursters. Together, these results demonstrate that, independent of how many preBötC neurons may be capable of intrinsic bursting, *I*_*NaP*_ can remain a critical component of the rhythmogenic mechanism beyond its contribution to network excitability.

To understand the potential interactions between *I*_*NaP*_ and synaptic dynamics for rhythm generation, we performed phase-specific manipulations of *I*_*NaP*_ activation/inactivation and synaptic activity. In control networks and following manipulations of spike shape (*g*_*SPK*_ = 15 *nS* or *g*_*AHP*_ = 35 *nS*) to eliminate intrinsic bursting, *I*_*NaP*_ inactivation and synaptic strength evolve with network firing rate along similar rotational trajectories during the respiratory cycle, comprised of burst initiation, burst termination, and the inter-burst interval (Fig. 3B). First, synaptic strength or *I*_*NaP*_ activation were manipulated at burst initiation (Fig. 3C), defined as the peak in *I*_*NaP*_ recovery from inactivation (*h*_*NaP*_). In all cases, when synapses were turned off at burst initiation, the expected network burst did not materialize, indicating that excitatory synaptic interactions are required to transition the network into bursts, even when all neurons are capable of intrinsic bursting. Similarly, if *I*_*NaP*_ activation (*m*_*NaP*_) was fixed in neurons that had not spiked yet at burst initiation, the network burst failed to occur under all conditions. Thus, with impaired *I*_*NaP*_ activation, synaptic interactions cannot initiate network bursts, and *vice versa*, illustrating that these can be interdependent properties for rhythm generation. Next, we characterized the role of synaptic depression and *I*_*NaP*_ inactivation in burst termination (Fig. 3D). In all three spike shape configurations (control, *g*_*SPK*_ = 15 *nS*, or *g*_*AHP*_ = 35 *nS*), synaptic depression was not essential for burst termination. However, without it burst duration was increased and the subsequent burst was delayed, particularly in the control network when all neurons were burst-capable. Interestingly, when *I*_*NaP*_ inactivation was fixed at burst initiation, network bursts only failed to terminate in control networks. In contrast, in networks with altered spike shape to eliminate intrinsic bursting, fixing *I*_*NaP*_ inactivation at burst initiation did not prevent burst termination, and only slightly increased burst duration and delayed the subsequent burst. Under these conditions, inter-burst intervals also became irregular (Fig. 3-Supplement 1), possibly indicative of a more stochastic process of burst initiation (Kam et al., 2013a,b; Feldman and Kam, 2015; Ashhad and Feldman, 2020; Ashhad et al., 2023). Finally, if synaptic depression and *I*_*NaP*_ inactivation were both fixed at burst initiation, bursts failed to terminate under all conditions. These results indicate that both *I*_*NaP*_ inactivation and synaptic depression can significantly contribute to, without being independently essential for, the termination of network bursts.

### preBötC rhythmogenesis is robust to partial *I*_*NaP*_ block

The effects of *I*_*NaP*_ antagonists on preBötC slice preparations have been inconsistent, fueling the debate surrounding the role of *I*_*NaP*_ in preBötC rhythm generation (see *Discussion*). Therefore, our model’s prediction that *I*_*NaP*_ is an essential element for preBötC rhythm-generation may seem controversial. This is due, in part, to the conflation of *I*_*NaP*_ with intrinsic bursting and the observation that *I*_*NaP*_-dependent intrinsic bursting is more sensitive to pharmacological manipulations than the network rhythm (Del Negro et al., 2002b, 2005; Phillips et al., 2022). To test this in our model network, we examined how rhythm generation and intrinsic bursting are affected by simulated attenuation of *I*_*NaP*_. Because the spike shape configurations described above represent the extreme scenarios (100% and 0% burst-capable) and spike heights of recorded preBötC neurons are generally higher and more variable than those produced by the model under control conditions, we simulated *I*_*NaP*_ blockade in networks where *g*_*SPK*_ was increased to 6 *nS* or uniformly distributed from 0− 12 *nS* (*g*_*SPK*_ = *U* (0, 12) *nS*), reducing the fraction of burst-capable neurons to 38% and 37%, respectively (Rekling and Feldman, 1998). Since effects of progressive *I*_*NaP*_ block were similar between spike shape configurations, simulations with *g*_*SPK*_ = *U* (0, 12) *nS, g*_*SPK*_ = 15 *nS, g*_*AHP*_ = 35 *nS*, and *g*_*SPK*_ = *g*_*AHP*_ = 0 *nS* are shown in (Fig. 4-Supplement 1 & 2). With *g*_*SPK*_ = 6 *nS* (Fig. 4A & B), progressive *I*_*NaP*_ blockade quickly reduced the fraction of burst-capable neurons and eliminated all intrinsic bursting when *g*_*NaP*_ was reduced by just ≈ 35% (Fig. 4C1-F1). Remarkably, much higher levels of *I*_*NaP*_ block were needed to prevent network rhythmogenesis, requiring *g*_*NaP*_ to be reduced by as much as 80 − 90% (compare white and red regions of Fig. 4E1). Furthermore, the sensitivity of the network rhythm was dependent on the excitability of the network prior to *I*_*NaP*_ block, with lower excitability networks being more sensitive to *I*_*NaP*_ block and higher excitability networks being less sensitive (compare points 1-5 in Fig. 4 E1 & F1). Notably, under either condition, once the rhythm was stopped by partial blockade of *I*_*NaP*_, an *I*_*NaP*_-dependent rhythm could be restored by increasing network excitability. Thus, these simulations illustrate how slightly different experimental conditions that influence preBötC excitability could lead to surprisingly variable results during pharmacological attenuation of *I*_*NaP*_ and different interpretations regarding its role in rhythm generation.

**Figure 4.**
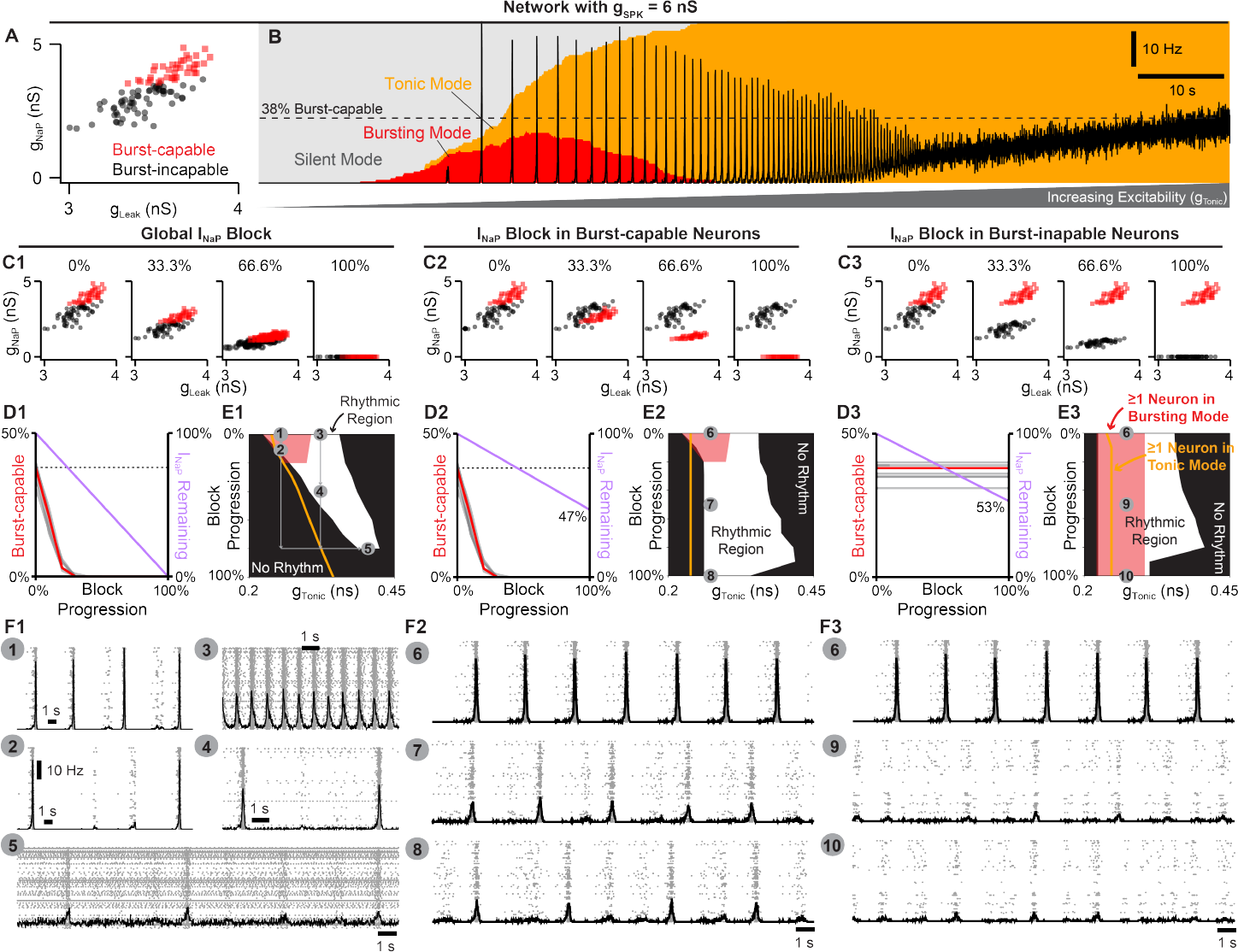
Selective block of *I*_*NaP*_ in burst-capable or burst-incapable neurons has similar consequences for rhythm generation. (A) Distributions of *g*_*NaP*_ and *g*_*Leak*_ among burst-capable (red) and incapable (black) neurons in a network with *g*_*SPK*_ = 6 *nS*. (B) Prevalence of silent, bursting, and tonic intrinsic cellular activities with overlaid network firing rate during increasing *g*_*Tonic*_ in the same network. (C1-3) Comparison of global *I*_*NaP*_ block (C1) vs. progressive *I*_*NaP*_ block specifically in neurons that are initially burst-capable (C2) or burst-incapable (C3). (D1-3) Fraction of the network that is burst-capable and amount of *I*_*NaP*_ remaining as a function of *I*_*NaP*_ block progression. (E1-3) Parameter space supporting intrinsic bursting (red) and network rhythmogenesis (white) as a function of excitability (*g*_*Tonic*_) during progressive *I*_*NaP*_ block. (F1-F3) Raster plots and overlaid network firing rate corresponding to points 1-10 shown in E1-3.

Because *I*_*NaP*_ is not specifically expressed in intrinsic bursting neurons making their selective manipulation experimentally intractable, we leveraged the advantages of computational modeling to compare how *I*_*NaP*_ in burst-capable and burst-incapable neurons contributes to network rhythmogenesis. This was done in model networks with *g*_*SPK*_ = 6 *nS* (Fig. 4 C2-F3) or *g*_*SPK*_ = *U* (0, 12) *nS* (Fig. 4-Supplement 2 C2-F3) by progressive suppression of *I*_*NaP*_ specifically in burst-capable neurons or burst-incapable neurons. Similar to global suppression of *g*_*NaP*_ (see Fig. 4 C1-F1), selective *I*_*NaP*_ suppression in burst-capable neurons (38% of the network) eliminated intrinsic bursting following a ≈ 35% reduction in *g*_*NaP*_. Yet, because only burst-capable neurons were affected, reducing *g*_*NaP*_ to 0 *nS* in this group of neurons only reduced the total *g*_*NaP*_ in the network by 47%. As a result, network rhythmogensis persisted despite the loss of intrinsic bursting and complete block of *I*_*NaP*_ in neurons that were initially burst-capable. On the other hand, selective suppression of *g*_*NaP*_ in burst-incapable neurons (62% of network) had no effect on the prevalence of intrinsic bursting, which remained constant at 38%, but led to a similar reduction in the total *g*_*NaP*_ in the network (53%). Notably, despite dramatically different effects on the prevalence of intrinsic bursting in the network, selective block of *I*_*NaP*_ in burst-capable or burst-incapable populations had surprisingly similar effects on network rhythmogenesis. Thus, in the model network, neurons with intrinsic bursting capabilities do not represent a functionally specialized neuronal population with a unique role in rhythm generation.

### Dynamic regulation of intrinsic bursting and pre-inspiratory spiking via small conditional modifications in spike shape

The manipulations of spike shape in the initial simulations (Figs. 1-4) were directly imposed. However, in neural systems, spike shape is dynamically regulated and can be altered indirectly by numerous conditional factors including e.g. temperature (Buzatu, 2009; Fohlmeister et al., 2010; Tang et al., 2010; Yu et al., 2012; Rinberg et al., 2013; Lujan et al., 2016; Tryba and Ramirez, 2004), oxygenation (Gruss et al., 2006), intracellular/extracellular ion concentrations (Strauss et al., 2008; Yang and Huang, 2022), and neurodevelopment (Ramoa and McCormick, 1994; Gao and Ziskind-Conhaim, 1998; Fry, 2006; Nakamura and Takahashi, 2007; Valiullina et al., 2016). Additionally, on shorter timescales, activity-dependent changes in spike height and AHP are common in neurons across the nervous system including preBötC neurons (Smith et al., 1991; Gray et al., 1999; Yamanishi et al., 2018) which may contribute to burst patterns and pre-inspiratory spiking (Abdulla et al., 2021). Intrinsic bursting in the preBötC seems to be affected by deliberate manipulations of some of these conditional factors (Del Negro et al., 2001; Mellen and Mishra, 2010; Peña et al., 2004; Tryba and Ramirez, 2004; Chevalier et al., 2016). Moreover, these factors also represent variables that are most likely to differ slightly between individual preBötC slice experiments and different research groups. Therefore, we explored whether indirect effects on spike shape during simulated changes in (1) oxygenation, (2) neurodevelopment, (3) extracellular potassium, and (4) temperature could capture experimental observations from preBötC slice preparations and provide conceptual insights into how conditional regulation of intrinsic bursting may obscure its perceived role in respiratory rhythm generation.

#### Hypoxia mediated changes in spike generation, intrinsic bursting, and network dynamics

When challenged acutely by exposure to hypoxia, the preBötC responds biphasically with augmented spiking activity and network burst frequency followed by suppressed activity and a gasping-like rhythm that appears more reliant on *I*_*NaP*_-dependent intrinsic bursting (Peña et al., 2004). Under hypoxic conditions, ATP production is decreased and the function of the Na+/K+-ATPase pump becomes impaired, disrupting ion gradients particularly via elevated intracellular sodium ([*Na*^+^]_*in*_) (Guatteo et al., 1998; Hellas and Andrew, 2021). As a result, spike-generating currents are weakened, and spike height and AHP are reduced (Gruss et al., 2006), which would be predicted to increase the prevalence of intrinsic bursting (see Fig. 1). However, if we consider that accumulation of [*Na*^+^]_*in*_ also reduces the driving force for *I*_*NaP*_, it becomes less clear how intrinsic bursting may be affected.

Therefore, we investigated how elevated [*Na*^+^]_*in*_ impacts spike shape, intrinsic bursting, and network dynamics. In the single-neuron model, increasing [*Na*^+^]_*in*_ decreased the sodium reversal potential (Fig. 5-Supplement 1A), which in turn reduced spike height and AHP (Fig. 5A & B). Increasing [*Na*^+^]_*in*_ also reduced neuronal excitability as indicated by higher levels of *g*_*Tonic*_ required to generate spiking/bursting (Fig. 5C). Despite reduced spike height and AHP, in model networks with distributed *g*_*SPK*_ (*U* (0, 12) *nS*), the percentage of burst-capable neurons was minimally affected and even decreased slightly with elevated [*Na*^+^]_*in*_ (Fig. 5D) due to the concurrent weakening of *I*_*NaP*_. However, in the neurons that remained burst-capable, intrinsic bursts became longer in duration with higher firing rates (Fig. 5D insets). In the synaptically coupled network, increasing [*Na*^+^]_*in*_ led to a decrease in the frequency and a small increase in the amplitude of network bursts before the rhythm was eventually lost at [*Na*^+^]_*in*_ *>* 21 *mM* (Fig. 5E).

**Figure 5.**
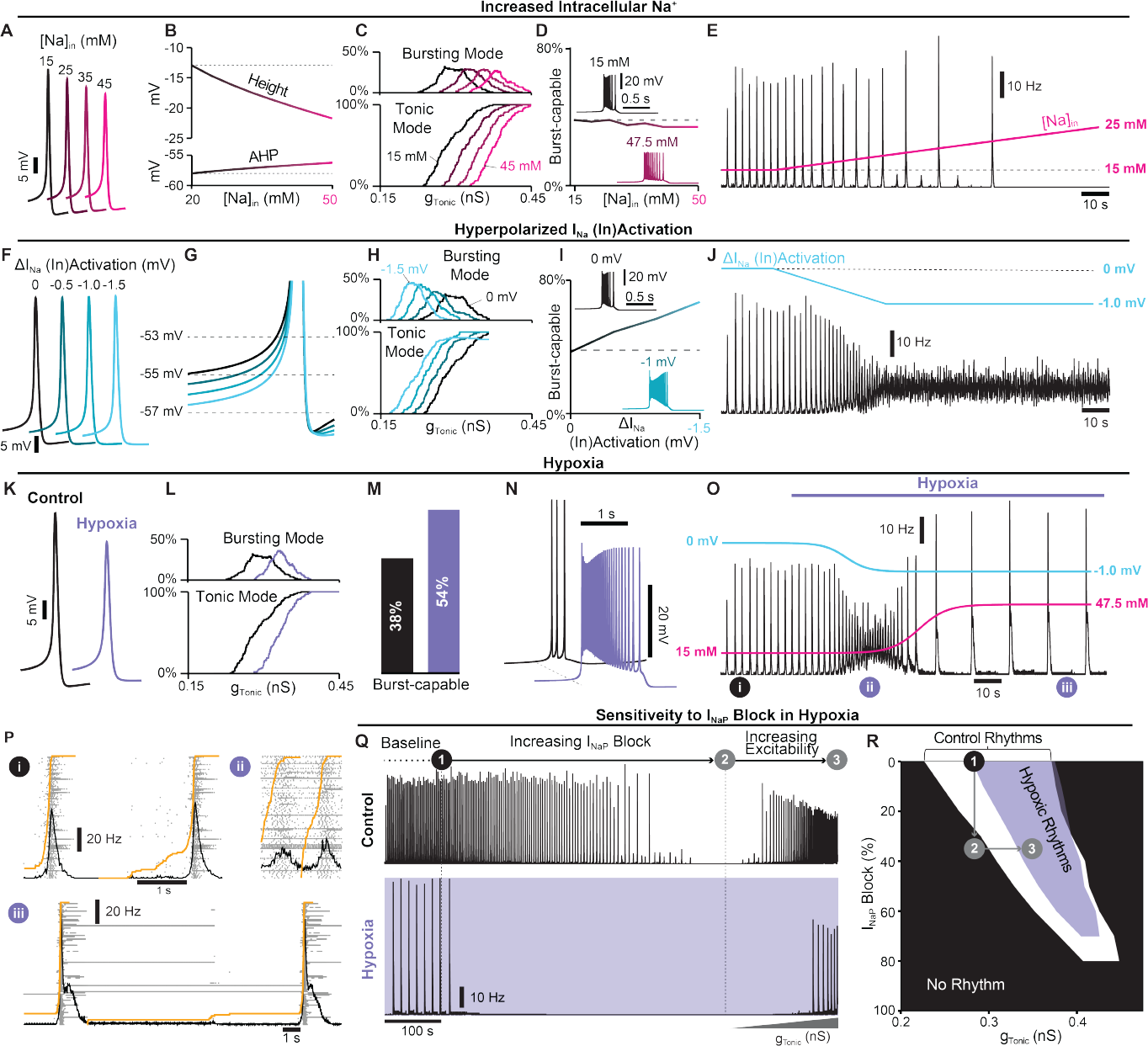
Simulated hypoxia alters spike generation, intrinsic bursting, network dynamics. (A) Example traces and (B) quantification of spike height and AHP during changes in [*Na*^+^]_*in*_. (C) Relationship between *g*_*Tonic*_ and the percentage of the network in tonic or bursting modes showing decreased excitability during elevated [*Na*^+^]_*in*_. (D) Number of burst-capable neurons in the network as a function of [*Na*^+^]_*in*_ with insets illustrating the impact on burst shape. (E) Effect of increasing [*Na*^+^]_*in*_ on the model network rhythm (*g*_*SPK*_ = *U* (0, 12)*nS*). (F) Example traces illustrating minimal changes in spike shape and (G) reduced spike threshold induced by a hyperpolarizing shift in the (in)activation dynamics of spike generating sodium currents (*I*_*Na*_ *& I*_*SPK*_). (H) Relationship between *g*_*Tonic*_ and the percentage of the network in tonic or bursting modes during *I*_*Na*_ *& I*_*SPK*_ (in)activation hyperpolarization. (I) Number of burst-capable neurons in the simulated preBötC network as a function of *I*_*Na*_ *& I*_*SPK*_ (in)activation hyperpolarization. Insets show representative intrinsic burster. (J) Network rhythm during linear hyperpolarization of *I*_*Na*_ *& I*_*SPK*_ (in)activation. (K) Example traces comparing spike shape under control and simulated steady-state hypoxia ([*Na*]_*in*_ = 47.5 *mM*, 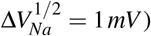). (L) Relationship between *g*_*Tonic*_ and the percentage of the population in tonic or bursting modes showing net decrease in excitability and (M) an increased percentage of the network that is burst-capable during hypoxia. (N) Effect of hypoxia on a representative intrinsic burster. (O) Network rhythm during simulated transition to hypoxia. (P) Example network traces before (i) and during the augmenting (ii) and gasping (iii) phases of the hypoxic response. (Q) Network activity and (R) parameter space supporting network rhythmogenesis during progressive *I*_*NaP*_ block under control (white) and hypoxic (purple) conditions.

Although increasing [*Na*^+^]_*in*_ revealed a rhythm that was similar to the gasp-like activity produced by the preBötC during hypoxia *in vitro*, it did not capture the typical biphasic response with an initial increase in network activity (Mironov et al., 1998; Thoby-Brisson and Ramirez, 2000; Peña et al., 2004; Garcia III et al., 2013). Recent studies suggest that the initial depolarization of neurons in response to hypoxia is due to relatively rapid (within 40 *s*) hyperpolarization of the voltage-dependent activation of fast spike-generating sodium channels. (Horn and Waldrop, 2000; Raley-Susman et al., 2001; Plant et al., 2016). Accordingly, we next tested how a hyperpolarizing shift of the voltage-dependent activation of 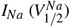 (Fig. 5-Supplement 1B) impacts spike-generation, intrinsic bursting, and network dynamics. Unexpectedly, in single neurons, we found that neither spike height nor AHP was significantly affected (Fig. 5F). However, the spike “threshold” was lowered (Fig. 5G) increasing neuronal excitability, as indicated by a leftward shift in the relationship between *g*_*Tonic*_ and the fraction of the network in tonic or bursting modes (Fig. 5H). Additionally, as 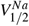 was hyperpolarized, the number of burst-capable neurons increased from 37% to 67% at 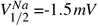 and burst duration increased with higher firing rates (Fig. 5I). In the synaptically coupled network, linearly hyperpolarizing 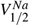 by 1 *mV* over 40 *s* led to an initial increase in network burst frequency followed by elimination of the rhythm (Fig. 5J).

Finally, we simulated the combined effects of altered 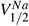 and elevated [*Na*^+^]_*in*_. In the single neuron model with [*Na*^+^]_*in*_ = 47.5 *mM* and 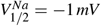, spike height and AHP were reduced by ≈ 7.5 *mV* and ≈ 1.4 *mV*, respectively (Fig. 5K) and excitability was reduced (Fig. 5L). In the model network, the fraction of burst-capable neurons increased from 38% to 54% (Fig. 5M) and the firing rate and duration of intrinsic bursts also increased (Fig. 5 N). Next, we simulated these consequences of hypoxia in the synaptically coupled network. Because the shift in 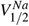 occurs relatively rapidly (Plant et al., 2016) and the resulting depolarization and increased spiking activity is expected to exacerbate [*Na*^+^]_*in*_ accumulation as the Na^+^/K^+^-ATPase pump becomes compromised, hypoxia was simulated as an initial change in 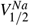 followed by accumulation of [*Na*^+^]_*in*_, each fit to a sigmoidal function. When combined, model networks responded with an initial increase in spiking activity and burst frequency followed by a rapid transition to a slow gasping-like rhythm, capturing the typical biphasic response of the preBötC to hypoxia (Fig. 5O). Specifically, simulated hypoxia resulted in the loss of pre-inspiratory spiking and transformation of burst shape from symmetrical to decrementing (Fig. 5P). Importantly, under these conditions, the network rhythm was also much more sensitive to *I*_*NaP*_ suppression (Fig. 5Q) as demonstrated for the preBötC network *in vitro* (Peña et al., 2004). However, this was not due to a change in the role of *I*_*NaP*_ or intrinsic bursting for rhythmogenesis, but was because of the reduced excitability that occurs with hypoxia. Consequently, similar to results under control conditions (see Fig. 4), if *I*_*NaP*_ was blocked by less than ≈ 70%, network rhythms in hypoxia could be restarted by increasing excitability (Fig. 5R).

#### Developmental changes in spike shape and intrinsic bursting mediated by increasing conductance densities

Experiments that have attempted to define the role of intrinsic bursting in preBötC rhythm generation have been restricted to prenatal or early postnatal development (Chevalier et al., 2016; Burgraff et al., 2022) with P0-P7 being the most common (Del Negro et al., 2002a; Pace et al., 2007b; Lorier et al., 2008; Ptak et al., 2005; Koizumi and Smith, 2008; Yamanishi et al., 2018). The possibility that intrinsic bursting may only be a feature of preBötC neurons during early development, while breathing must continue throughout life, has been a longstanding criticism of the *pacemaker hypothesis*. In general, during embryonic and postnatal development, spike height and AHP increase, while spike duration decreases (Ramoa and McCormick, 1994; Gao and Ziskind-Conhaim, 1998; Fry, 2006; Nakamura and Takahashi, 2007; Valiullina et al., 2016). These changes are primarily due to increasing densities of voltage-gated ion channels (Huguenard et al., 1988; Gao and Ziskind-Conhaim, 1998; Fry, 2006; Valiullina et al., 2016). Thus, we performed simulations during scaling of ionic conductances to predict how neurodevelopment may affect intrinsic bursting and network dynamics via changes in spike shape and *I*_*NaP*_ conductances densities.

In single model neurons, we simulated changes in voltage-gated conductance densities by applying a scaling factor ranging from 0.25*X* to 2.5*X* to all voltage-gated conductances except *g*_*NaP*_ (Fig. 6A). When conductances were scaled down, spike height and AHP were reduced and spike duration became longer. Notably, in model neurons, further reducing spike height and AHP by down-scaling conductances rendered them unable to generate sustained trains of spikes (Fig. 6B), as observed in early embryonic development (Gao and Ziskind-Conhaim, 1998; Boeri et al., 2021). Conversely, as conductance densities were scaled up, spike height and AHP increased, and spike duration became shorter (Fig. 6B & C). Conductance scaling also reduced cellular excitability as indicated by higher values of *g*_*Tonic*_ required to initiate bursting or tonic spiking (Fig. 6G). Among burst-capable neurons, simulation of neurodevelopment via conductance scaling transformed the shape of intrinsic bursts, which resembled long-duration plateau-like bursters with down-scaling (Chevalier et al., 2016) and became shorter in duration with up-scaling until neurons transitioned to tonic spiking and were rendered burst-incapable (Fig. 6F). Interestingly, the frequency range of these plateau-like bursters is very restricted (≈ 0.05 *Hz* to ≈ 0.1 *Hz*) and their bursting capabilities are highly insensitive to *I*_*NaP*_ attenuation ((Fig. 6 Supplement 2A,B), consistent with experimental recordings (Chevalier et al., 2016).

**Figure 6.**
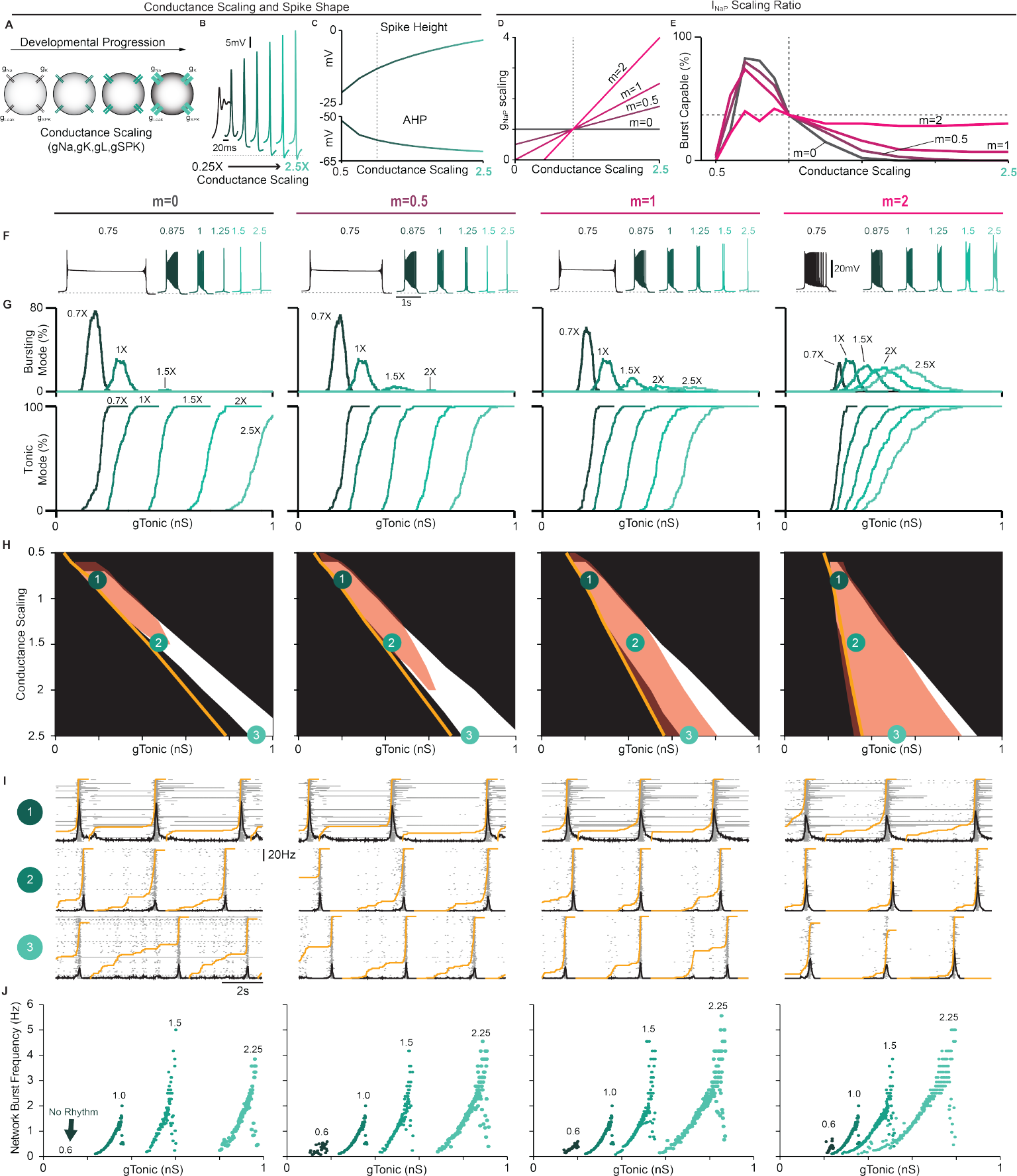
Predicted developmental changes in spike shape, intrinsic bursting and network dynamics due to changing conductance densities. (A) Illustration of conductance scaling during development. (B) Example traces and (C) quantification of spike height and AHP as conductances are scaled. (D) Ratios of concurrent *g*_*NaP*_ scaling (*m* = 0 − 2) and (E) percentage of the network (*g*_*SPK*_ = *U* (0, 12) *nS*) that is burst-capable as conductances are up- or down-scaled from control values (dashed vertical line). (F) Example intrinsic bursting neurons during conductance scaling with *m* = 0 − 2. (G) Decreased excitability with conductance scaling as indicated by a rightward shift in the level of *g*_*Tonic*_ needed to initiate intrinsic bursting or tonic spiking. (H) Comparison of parameter space that supports intrinsic bursting (red) and network rhythmogenesis (white) as conductances are scaled with *m* ranging from 0− 2 (Orange lines indicate *g*_*Tonic*_ where ≥1 neuron enters tonic spiking mode). (I) Raster plots and overlaid network firing rate corresponding to points 1-3 in (H) (Orange line indicate the percentage of neurons active since the preceding network burst). (J) Relationship between excitability (*g*_*Tonic*_) and network burst frequency as conductances are scaled with *m* ranging from 0 − 2.

To examine how concurrent neurodevelopmental changes in *g*_*NaP*_ may affect intrinsic bursting and network dynamics, we added scaling to *g*_*NaP*_ ranging from 0*X* to 2*X* the scaling factor applied to other voltage-gated conductances (*m* = 0 − 2, Fig 6D). In model networks (*g*_*SPK*_ = *U* (0, 12) *nS*), we quantified the proportion of burst-capable neurons as conductance densities undergo scaling with varied ratios of concurrent *g*_*NaP*_ scaling (Fig.6E). Similar simulations in networks with *g*_*SPK*_ = 0, 6, or 12 are shown in Fig. 6-Supplement 1. In all cases, when conductances were low (scaling factor *<* 0.5*X*), intrinsic bursting was not possible in any neurons. When *g*_*NaP*_ was concurrently scaled at 0, 0.5, or 1*X* the scaling factor for other conductances (*m* = 0, 0.5, or 1), the fraction of burst-capable neurons quickly increased with up-scaling, reaching a peak of ≈ 70 − 80% at a scaling factor of 0.75*X*, and then declining to 38% under control conditions (scaling factor = 1*X*). As conductance densities were further scaled up, the number of burst-capable neurons continued to decline until intrinsic bursting was lost or only possible in a small fraction of the population. When the ratio of *g*_*NaP*_ scaling was 2*X* (*m* = 2), the peak in burst-capable neurons at scaling *<* 1*X* was diminished, but more neurons retained bursting capabilities as scaling increased (Fig. 6E-H). This decline in intrinsic bursting typically corresponded with increasing pre-inspiratory spiking activity (Fig. 6I), and also expanded the excitability (*g*_*Tonic*_) range that supported rhythmogenesis, allowing the network to produce a much wider range of frequencies (Fig. 6J). In all cases, rhythmogenesis remained dependent on *I*_*NaP*_ and, interestingly, intrinsic bursting became more sensitive to *I*_*NaP*_ attenuation whereas the sensitivity of network rhythmogenesis was unchanged or slightly decreased (Fig. 6-Supplement 2C). Collectively, these results illustrate how developmental factors that affect spike shape may give rise to changes in the prevalence of intrinsic bursting. These results also illustrate that, even with scaling up of *g*_*NaP*_, intrinsic bursting can remain a feature limited to a subset of neurons within a certain developmental period, supporting the interpretation that intrinsic bursting is a side effect of *I*_*NaP*_-dependent rhythm generation without a specialized functional role.

#### In vitro to in vivo: impact of extracellular potassium and temperature on cellular and network bursting

Inherent in the process of creating the *in vitro* preBötC slice, excitatory inputs from regions outside the preBötC that regulate its activity are removed. To compensate for this loss of excitability, elevating the concentration of potassium in the bathing solution to between 8 *mM* and 9 *mM* is standard practice to promote reliable rhythmic activity from the preBötC. In addition, slices are typically maintained at a subphysiological temperature (27^*°*^*C*) to extend the viability of the preparation (Smith et al., 1991; Funk et al., 1993; Del Negro et al., 2002a; Koizumi and Smith, 2008; Smith et al., 2007; Yamanishi et al., 2018). The possibility that these artificial conditions may also impact intrinsic bursting has been an enduring criticism of the *pacemaker hypothesis* and the *in vitro* preparation in general. Indeed, to what extent the biophysical mechanisms of rhythm generation seen under *in vitro* conditions are representative of normal physiology remains unclear and has been an important caveat common to the study of CPGs in general (MacKay-Lyons, 2002; Grillner et al., 2005; Feldman and Kam, 2015; Marder and Bucher, 2001; Marder et al., 2005). Because spike shape changes with both temperature (Buzatu, 2009; Fohlmeister et al., 2010; Tang et al., 2010; Yu et al., 2012; Rinberg et al., 2013; Lujan et al., 2016) and extracellular potassium ([*K*^+^]_*ext*_) (Strauss et al., 2008; Yang and Huang, 2022), we explored how these variables may impact intrinsic bursting and network rhythms to better understand whether the essential biophysical mechanisms of rhythm generation are conserved under simulated temperatures and [*K*^+^]_*ext*_ associated with *in vitro* and *in vivo* conditions.

In single model neurons, reducing [*K*^+^]_*ext*_ (represented by the parameter *K*_*Bath*_) hyperpolarized the *K*^+^ and leak reversal potentials (*E*_*K*_ and *E*_*Leak*_, Fig. 7-Supplement 1A), which increased the spike AHP and slightly reduced spike height (Fig. 7A and B), as expected (Strauss et al., 2008; Yang and Huang, 2022; Bacak et al., 2016; Abdulla et al., 2021; Powell and Brown, 2021). In the model preBötC network with distributed spike heights (*g*_*SPK*_ = *U* (0, 12) *nS*), decreasing [*K*^+^]_*ext*_ from a baseline value of 8.5 *mM* reduced excitability as indicated by an increased *g*_*Tonic*_ required for neurons to enter spiking/bursting modes (Fig. 7C). Additionally, decreasing [*K*^+^]_*ext*_ quickly reduced and then eliminated burst-capable neurons at [*K*^+^]_*ext*_ *<* 5 *mM* (Fig. 7D), consistent with experimental observations (Del Negro et al., 2001; Mellen and Mishra, 2010; Johnson et al., 2016). Reducing [*K*^+^]_*ext*_ below 5 *mM* also led to the cessation of the network rhythm (Fig. 7E), as is typical in most *in vitro* preBötC slice preparations (Smith et al., 1991; Funk et al., 1993; Del Negro et al., 2001; Kallurkar et al., 2020). However, if a source of excitatory drive was provided to the network to increase its excitability (*g*_*Tonic*_), as expected to be present *in vivo*, the network rhythm re-emerged despite the continued absence of intrinsic bursting (Fig. 7E). It is also notable that, at [*K*^+^]_*ext*_ = 4 *mM*, the onset of simulated hypoxia (see Fig. 5 also revealed a transient rhythm that re-emerged and persisted following recovery from hypoxia (Fig. 7-Supplement 2A), as has been observed experimentally (Mironov, 2013). In the model, this is due to short-term [*Na*^+^]_*in*_ dynamics and the long-lasting hyperpolerizing shift in the voltage-dependence of *I*_*Na*_ (in)activation.

**Figure 7.**
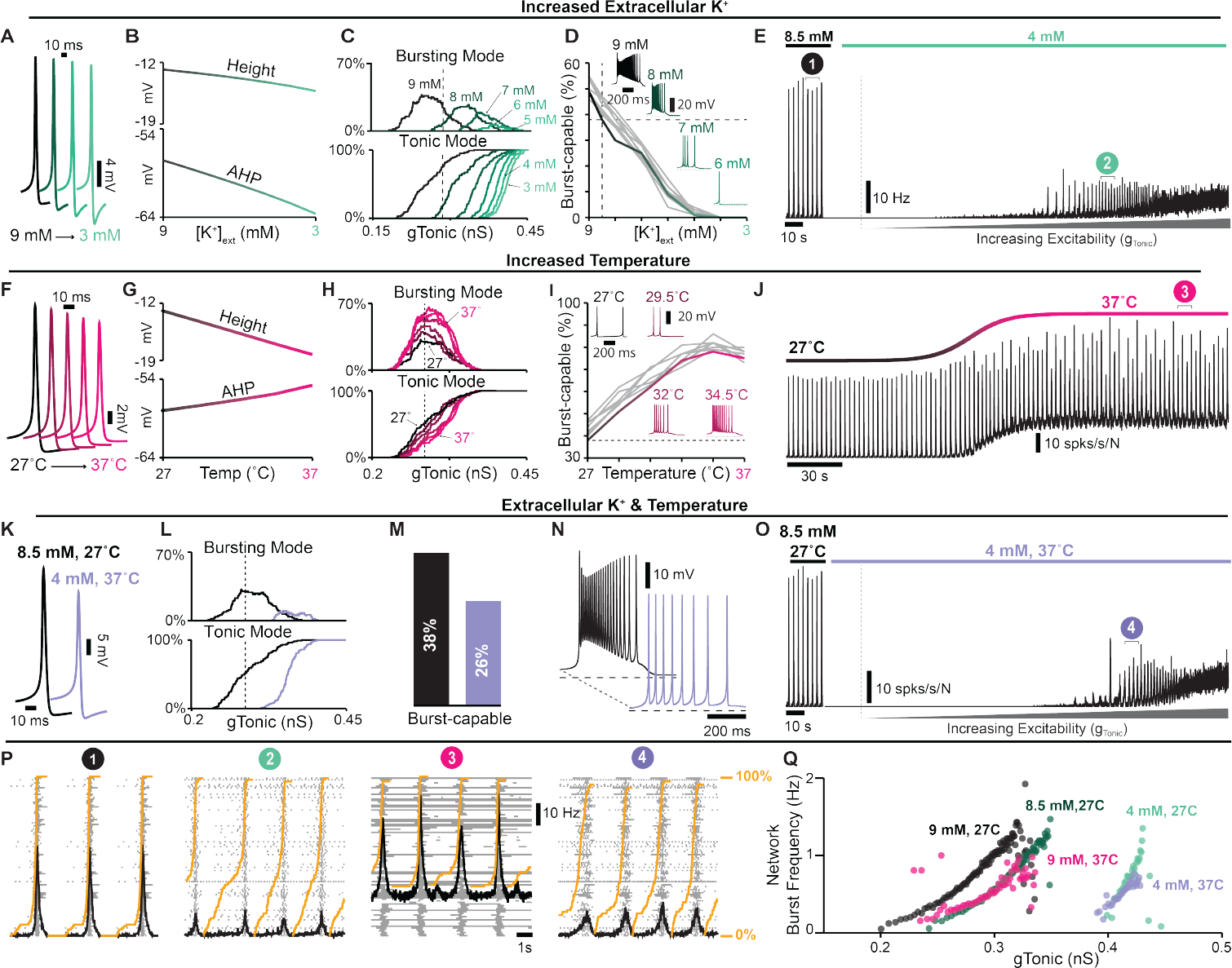
Regulation of spike shape, intrinsic bursting and network dynamics by 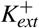 concentrations and temperatures associated with *in vitro* and *in vivo* conditions. (A) Example traces and (B) quantified spike height and AHP during changes in [*K*^+^]_*ext*_. (C) Relationship between *g*_*Tonic*_ and the percentage of the network in bursting or tonic modes showing reduced excitability at lower [*K*^+^]_*ext*_. (D) Percentage of burst-capable neurons in the network as a function of [*K*^+^]_*ext*_ with insets of a representative intrinsic burster. (E) Network rhythm at 8.5*mM* and 4*mM* [*K*^+^]_*ext*_ during increasing *g*_*Tonic*_. (F) Example traces and (G) quantified spike height and AHP during changes in temperature. (H) Relationship between *g*_*Tonic*_ and the percentage of neurons in busting or tonic modes. (I) Percentage of burst-capable neurons as a function of temperature with insets showing representative intrinsic burster. (J) Network rhythm during an increase in temperature from 27^*°*^*C* to 37^*°*^*C*. (K) Example spike shapes under *in vitro* ([*K*^+^]_*ext*_ = 8.5 *mM, T* = 27^*°*^*C*) and *in vivo*-like ([*K*^+^]_*ext*_ = 4 *mM, T* = 37^*°*^*C*) conditions. (L) Net decrease in excitability, indicated by a rightward shift in the relationship between *g*_*Tonic*_ and the percentage of the population in bursting or tonic modes, and (M) the percentage of burst-capable neurons under *in vivo*-like conditions. (N) Representative intrinsic burster in each condition. (O) Network rhythm during transition from *in vitro* to *in vivo* [*K*^+^]_*ext*_ and temperature and during increasing excitatory drive (*g*_*Tonic*_). (P) Rasters and overlaid population firing rate for points i-iv shown in E, J, and O (Orange lines indicate fraction of the network active since preceding burst). (Q) Effects of [*K*^+^]_*ext*_ and/or temperature on the relationship between excitability and network burst frequency.

Next, we considered the potential consequences of temperature. Spike height and AHP are known to decrease with increasing temperature (Buzatu, 2009; Fohlmeister et al., 2010; Tang et al., 2010; Yu et al., 2012; Rinberg et al., 2013; Lujan et al., 2016) including in preBötC neurons (Tryba and Ramirez, 2004). These changes are thought to be largely due to faster dynamics of voltage-gated channels and increased neuronal capacitance (Matteson and Armstrong, 1982; Collins and Rojas, 1982; Fohlmeister et al., 2010; Yu et al., 2012; Shapiro et al., 2012; Pinto et al., 2021, 2022; Plaksin et al., 2018). Therefore, to simulate changes in temperature, we added temperature dependence to voltage-dependent rate constants and cell capacitance such that all rate constants are reduced by ≈ 70% and capacitance increases by ≈ 1 *p f* over the 10^*°*^C change from 27^*°*^C to 37^*°*^C (Fig. 7-Supplement 1B & C), see *Materials and Methods* for a full description. With these dependencies, simulating an increase in temperature decreased spike height and AHP (see Fig. 7F), consistent with experimental observations (Buzatu, 2009; Fohlmeister et al., 2010; Tang et al., 2010; Yu et al., 2012; Rinberg et al., 2013; Lujan et al., 2016). In the network, increasing temperature increased the possible number of neurons concurrently in a bursting mode but did not impact excitability as indicated by an unchanged *g*_*Tonic*_ required to depolarize neurons into spiking/bursting modes (Fig. 7H). Importantly, the number of burst-capable neurons was increased from 38% at 27^*°*^C to 75% at 37^*°*^C (Fig. 7I) and interestingly, some neurons that were initially in a bursting mode transitioned to tonic spiking mode and vice versa, consistent with the findings of Tryba and Ramirez (2004) (Fig. 7-Supplement 1D). In the synaptically coupled network, this resulted in a shift in the baseline spiking activity of the network and an increase in burst frequency (Fig. 7J), as observed experimentally (Tryba and Ramirez, 2003, 2004).

Finally, we examined how differences in [*K*^+^]_*ext*_ and temperature may impact the cellular- and network-level properties of the preBötC *in vitro* and *in vivo*. In single model neurons, simultaneously decreasing [*K*^+^]_*ext*_ from 8.5 to 4 *mM* and increasing temperature from 27^*°*^C to 37^*°*^C resulted in a net decrease in spike height and AHP (Fig. 7K). In the network, this change in [*K*^+^]_*ext*_ and temperature reduced excitability, requiring higher *g*_*Tonic*_ for neurons to enter tonic/bursting modes, decreased the possible number of neurons concurrently in bursting mode (Fig. 7L), and reduced the number of burst-capable neurons from 38% to 26% (Fig. 7M). Despite the persistence of intrinsic bursting capabilities, this change in [*K*^+^]_*ext*_ and temperature caused cessation of the network rhythm due to reduced excitability. Accordingly, if an excitatory input was applied to the network (*g*_*Tonic*_) the network rhythm re-emerged (Fig. 7O). Under these conditions, the dynamic frequency range of the network remained largely unchanged or slightly reduced (see Fig. 7O), and there was a higher fraction of the network participating in pre-inspiratory activity (Fig. 7P). Interestingly, in the model, if synaptic strength was increased at physiological [*K*^+^]_*ext*_, as suggested by prior experiments (Rimmele et al., 2017; Rausche et al., 1990; Czéh et al., 1988; Gonzalez et al., 2022; Erulkar and Weight, 1977), the network rhythm increased in amplitude and became more robust (Fig. 7-Supplement E). Despite the differences in cellular activity phenotypes (intrinsic bursting and pre-inspiratory spiking) and network activity between simulated [*K*^+^]_*ext*_ and temperature conditions *in vitro* and *in vivo*, rhythmogenesis remained dependent on *I*_*NaP*_ and excitatory synaptic connections under all conditions (Fig. 7-Supplement F).

## DISCUSSION

In this study, we address the longstanding debate surrounding respiratory rhythm generation using computational modeling to disentangle the conflated role(s) of *I*_*NaP*_ and intrinsic bursting. By characterizing how the voltage-dependent properties of *I*_*NaP*_ interact with spike shape, we discover that small changes in spike height and/or AHP can transition intrinsic bursting neurons to tonic spiking and render them incapable of intrinsic bursting (Fig. 1). In an established preBötC network model that is commonly viewed as a quantitative realization of the *pacemaker hypothesis*, we leverage this interaction to selectively render all neurons capable or incapable of intrinsic bursting. By doing so, we demonstrate that preBötC rhythmogenesis persists in both extremes - when intrinsic bursting is not possible and neurons exhibit intrinsically tonic activity associated with pre-inspiratory spiking in the network, and also when all neurons are capable of intrinsic bursting but the network lacks pre-inspiratory spiking. (Fig. 2). Thus, while these phenotypes of preBötC neurons may be present, they are conditional and do not represent essential rhythmogenic elements of the network. Instead, regardless of the amount of intrinsic bursting or pre-inspiratory spiking, rhythmogenesis *per se* remains dependent on interactions between *I*_*NaP*_ and recurrent synaptic excitation (Fig. 3-4). Consistent with these findings and extensive experimental observations often cited in support of either the *pacemaker hypothesis* or *burstlet theory*, we illustrate how conditional factors that impact spike shape including oxygenation (Fig. 5), development (Fig. 6), extracellular potassium, and temperature (Fig. 7) can substantially alter the relative abundance of intrinsic bursting and pre-inspiratory spiking without precluding rhythm generation. Thus, rather than being rhythmogenic, we propose that such changes in the activity patterns of preBötC neurons are consequences of a network that evolved to be robust, ensuring breathing persists despite developmental or environmental changes while remaining able to accommodate the wide range of breathing patterns associated with its physiological, behavioral, and emotional integration.

Over the last three decades, impressive progress has been made toward understanding the central control of breathing (Ramirez and Baertsch, 2018; Molkov et al., 2017; Del Negro et al., 2018; Ashhad et al., 2022; Guyenet and Bayliss, 2015; Feldman et al., 2003). However, the debate surrounding preBötC rhythm generation has remained largely unchanged. This stems, in part, from the simplistic framing of the *pacemaker* and *group-pacemaker/burstlet theories*. Although attractive and broadly accessible, this simplicity supports a false dichotomy that obscures more nuanced interpretations critical to achieve a consensus view.

First, it implies that these theories are mutually exclusive. For example, in *group-pacemaker*-based interpretations, rhythmogenesis is described as an ‘emergent’ property of the network because it arises from interactions among “non-rhythmic” intrinsically tonic neurons. On the other hand, in networks that contain intrinsically bursting neurons, rhythmicity is often assumed to be driven by the activity of these bursting neurons as if they were “pacing” the network. However, intrinsic burst frequencies among pacemaker neurons are quite variable (Johnson et al., 1994) and, as with any other preBötC neuron, intrinsic bursting neurons are embedded within a recurrently connected network. Therefore, synaptic interactions are *always* required to coordinate cellular activity into a coherent network rhythm regardless of the intrinsic spiking patterns of its constituent neurons (see Fig. 3). Thus, in our view, all network rhythms are ‘emergent’ as they arise from the collective activity of individual neurons coordinated via synaptic interactions. Moreover, tonic spiking and intrinsic bursting are both rhythmic processes, and therefore the synchronization of individual spikes or clusters of spikes (i.e. bursts) across the network via synaptic interactions may share far more similarities than differences. Indeed, in the model presented here, we find that the initiation of network bursts depends on both *I*_*NaP*_ activation and recurrent excitatory interactions, whereas *I*_*NaP*_ inactivation and synaptic depression both contribute to burst termination (Fig. 3). Thus, the roles of network interactions and the intrinsic properties of the neurons within it should not be considered separable or “one or the other”. Instead, we propose that “the” mechanism of rhythm generation involves multiple interacting and interdependent properties of the preBötC.

Second, with this framing, *I*_*NaP*_ has become misconstrued with intrinsic bursting and the *pacemaker hypothesis*. This may reflect, in part, the way in which intrinsic bursting and *I*_*NaP*_ were initially characterized in the preBötC-first with the discovery of pacemaker neurons (Smith et al., 1991), followed by incorporation of *I*_*NaP*_ into computational models with the goal of replicating the pacemaker phenotype (Butera et al., 1999a), and finally subsequent experimental confirmation of *I*_*NaP*_ expression and *I*_*NaP*_-dependent pacemaker neurons in the preBötC (Del Negro et al., 2002a; Koizumi and Smith, 2008). This progression of discovery strongly supported the assumption that the purpose of *I*_*NaP*_ in the preBötC is to endow some neurons with intrinsic bursting properties, which in turn drives rhythmic activity of the network. However, rather than a driver of rhythm, our findings suggest that it may be more accurate to view intrinsic bursting as a consequence of *I*_*NaP*_-dependent rhythm generation that is only possible in neurons that happen to have a certain combination of properties including, but not limited to, *g*_*NaP*_, *g*_*Leak*_, and any of the many properties that influence spike shape (Buzatu, 2009; Fohlmeister et al., 2010; Tang et al., 2010; Yu et al., 2012; Rinberg et al., 2013; Lujan et al., 2016; Tryba and Ramirez, 2004; Gruss et al., 2006; Strauss et al., 2008; Yang and Huang, 2022; Ramoa and McCormick, 1994; Gao and Ziskind-Conhaim, 1998; Fry, 2006; Nakamura and Takahashi, 2007; Valiullina et al., 2016; Strauss et al., 2008), see Figs. 2–7. Taking this view, one need not consider the small subset of neurons that are capable of intrinsic bursting to be a specialized cell type with a unique functional role. Indeed, our simulations demonstrating that blockade of *I*_*NaP*_ specifically in burst-capable or incapable neurons has similar consequences for network rhythmogenesis (Fig. 4) support this view. This interpretation is also supported by experimental observations that the preBötC rhythm can persist even after intrinsic bursting is apparently abolished by pharmacological or genetic attenuation of *I*_*NaP*_ (Del Negro et al., 2002b; da Silva et al., 2019). Due to the conflation of *I*_*NaP*_ and intrinsic bursting, these findings have reinforced the idea that *I*_*NaP*_ is not obligatory for preBötC rhythm generation. However, the role of *I*_*NaP*_ in rhythm generation need not be restricted to intrinsic bursting. Indeed, our simulations clearly illustrate that *I*_*NaP*_ can be critical for rhythm generation independent of any requirement for intrinsic bursting neurons (see Fig. 2,3), and that *I*_*NaP*_-dependent preBötC rhythms can persist after intrinsic bursting is abolished following partial *I*_*NaP*_ block (see Fig. 4, 5, 6-Supplement 2, and 7-Supplement 1). These simulations are an important proof of concept that equating *I*_*NaP*_ and intrinsic bursting is an oversimplification.

The debate surrounding the role of *I*_*NaP*_ has also been exacerbated by the seemingly inconsistent effects of *I*_*NaP*_ blockers. (Del Negro et al., 2002a; Peña et al., 2004; Ramirez et al., 2004; Del Negro et al., 2005; Feldman and Del Negro, 2006; Smith et al., 2007; Pace et al., 2007b; Koizumi and Smith, 2008; Kam et al., 2013a,b; Ashhad and Feldman, 2020). For example, in cases where *I*_*NaP*_ blockers have failed to eliminate the preBötC rhythm, proponents of the *pacemaker hypothesis* often contend that this is because block of *I*_*NaP*_ was incomplete due to e.g. insufficient diffusion of drug into the tissue or too low of a dose used (Koizumi and Smith, 2008; Phillips and Rubin, 2019; Phillips et al., 2022). On the other hand, in cases where *I*_*NaP*_ blockers have eliminated the preBötC rhythm, proponents of *group pacemaker/burstlet theory*, generally attribute this to *I*_*NaP*_ ‘s contribution to cellular excitability rather than an essential role in rhythmogenesis *per se* (Pace et al., 2007b). This is supported by some experimental observations that, following elimination of the preBötC rhythm with the *I*_*NaP*_ blocker Riluzole, rhythmicity could be restored by application of substance P to increase preBötC excitability (Pace et al., 2007b). Here, we illustrate how *I*_*NaP*_’s contribution to preBötC excitability can be a key factor underlying the widely variable responses of the preBötC rhythm to suppression of *I*_*NaP*_ (Fig. 4), providing a plausible explanation for these apparently discrepant findings. Because sufficient cellular excitability is also critical for preBötC rhythmogenesis, if the level of excitability is initially low, a modest suppression of *I*_*NaP*_ (*<*≈ 10%) will quickly stop the rhythm because the total excitability becomes insufficient for rhythmogenesis. However, if baseline excitability is higher, it becomes much more difficult for *I*_*NaP*_ suppression to eliminate rhythm generation (Fig. 4). Further, once the rhythm has been stopped by partial suppression of *I*_*NaP*_, it can be restarted by increasing excitability so long as *I*_*NaP*_ has not been suppressed by more than 80 − 85% (Fig. 4), consistent with experiments suggesting that the preBötC rhythm can only be restarted with substance P when *I*_*NaP*_ block is incomplete (Koizumi and Smith, 2008). Thus, the wide variation in the amount of *I*_*NaP*_ suppression required to stop rhythm generation does not indicate that *I*_*NaP*_ has a more important rhythmogenic role in one condition vs. another. Nor does the ability to recover rhythmicity by increasing excitability suggest that *I*_*NaP*_ is not an essential element of rhythmogenesis. Instead, independent from its contribution to excitability, rhythm generation remains dependent on *I*_*NaP*_ due to its contribution to burst initiation (Fig. 3), which requires substantial attenuation of *I*_*NaP*_ (*>* 80 − 85%) to be impaired. This is consistent with optogenetic manipulations of preBötC excitability during graded pharmacological block of *I*_*NaP*_ (Phillips et al., 2022). Notably, the prevalence of burst-capable neurons in the network has little effect on the relationship between excitability and the sensitivity of the rhythm to *I*_*NaP*_ suppression (Fig. 4-Supplement 1 & 2). Collectively, these simulations illustrate that *I*_*NaP*_ ‘s role in preBötC rhythm generation is not limited to its contribution to excitability or intrinsic bursting, and provide important conceptual insight into why experimental efforts to define the role of *I*_*NaP*_ in preBötC rhythm generation have been inconsistent and difficult to interpret.

Third, both theories generally overlook the conditional nature of intrinsic bursting and pre-inspiratory spiking (Feldman and Del Negro, 2006; Feldman et al., 2013; Del Negro et al., 2018; Molkov et al., 2017; Ramirez and Baertsch, 2018; Smith et al., 2000; Smith, 2022). Importantly, our simulations reveal an inverse relationship between the prevalence of intrinsic bursting and pre-inspiratory spiking in the network that can be profoundly altered by small changes in spike shape. Conditions that may influence this balance can be artificial or physiological. Indeed, a long-standing critique of the *pacemaker hypothesis* (and by association *I*_*NaP*_) is the lack of evidence for intrinsic bursting in adult animals *in vivo*. Although the absence of evidence is not evidence of absence, this suggests that intrinsic bursting could be 1) restricted to early development and/or 2) an artifact of the artificial conditions used to record from *in vitro* slice preparations. Many of these artificial and physiological factors can affect spike shape (Buzatu, 2009; Fohlmeister et al., 2010; Tang et al., 2010; Yu et al., 2012; Rinberg et al., 2013; Burgraff et al., 2022; Abdulla et al., 2021; Lujan et al., 2016; Tryba and Ramirez, 2004; Strauss et al., 2008; Yang and Huang, 2022; Ramoa and McCormick, 1994; Gao and Ziskind-Conhaim, 1998; Fry, 2006; Nakamura and Takahashi, 2007; Valiullina et al., 2016; Ptak et al., 2005; Krey et al., 2010; Phillips et al., 2018; Revill et al., 2021), and would therefore be expected to shift the prevalence of intrinsic bursting and pre-inspiratory spiking without altering the role of *I*_*NaP*_ in rhythm generation.

*In vitro* and *in vivo* experiments are typically performed at different [*K*^+^]_*ext*_ and temperature. In preBötC slices, artificially increasing [*K*^+^]_*ext*_ promotes rhythmogenesis and also intrinsic bursting (Del Negro et al., 2001; Mellen and Mishra, 2010). In contrast, lower [*K*^+^]_*ext*_ (sometimes with altered [Ca^2+^]) promotes weaker “burstlet” rhythms hypothesized to be driven by pre-inspiratory spiking rather than intrinsic bursting or *I*_*NaP*_ (Kam et al., 2013a,b; Feldman and Kam, 2015). Consistent with these experimental observations, lowering [*K*^+^]_*ext*_ in the model reduces the number of burst-capable neurons in the network due to an increase in spike AHP. Experimentally, at physiological [*K*^+^]_*ext*_ (Zacchia et al., 2016; Takahashi et al., 1981; Okada et al., 2005), intrinsic bursting is eliminated and the network rhythm stops (Del Negro et al., 2001). However, the latter is a consequence of reduced cellular excitability at lower [*K*^+^]_*ext*_ because rhythmogenesis can be restored if excitatory drive is increased (Fig. 7), as expected *in vivo* due to the presence of e.g. neuromodulatory and chemoreceptor inputs (Souza et al., 2023). Under these conditions, intrinsic bursting remains absent, pre-inspiratory spiking is increased, and characteristics of the network rhythm become more consistent with “burstlets” (Fig. 7). However, another artificial aspect of *in vitro* experiments is low temperature, which has an inverse relationship with spike height and AHP (Strauss et al., 2008; Yang and Huang, 2022); Fig. 7. Warmer temperatures *in vivo* are therefore expected to counteract the effects of low [*K*^+^]_*ext*_ on intrinsic bursting. As a result, our model predicts that at physiological temperature and [*K*^+^]_*ext*_ some neurons remain burst-capable, consistent with experiments that have identified intrinsic bursting preBötC neurons at physiological [*K*^+^]_*ext*_ but at slightly warmer temperatures (30^*°*^*C* - 31^*°*^*C*) (Tryba et al., 2003; St.-John et al., 2009).

*In vitro* and *in vivo* experiments are generally performed at different stages of development. Intrinsic bursting in the preBötC is often thought to be most prevalent during early development (Hilaire and Duron, 1999; Smith et al., 2000), and attempts to record rhythmic preBötC activity in slices from rodents *>*≈ *P*14 have been generally unsuccessful with few exceptions (Ramirez et al., 1996). This has further reinforced the assumptions that intrinsic bursting drives rhythmic activity of the preBötC *in vitro*, and that the preBötC rhythm *in vivo* must be generated by a distinct mechanism such as reciprocal inhibition (Smith et al., 2000; Richter and Smith, 2014). Our simulations illustrate how the abundance of burst-capable neurons can peak during early development due to changes in spike shape that are expected as the densities of voltage-gated conductances increase (Ramoa and McCormick, 1994; Gao and Ziskind-Conhaim, 1998; Fry, 2006; Nakamura and Takahashi, 2007; Valiullina et al., 2016); Fig. 6. However, as discussed above, this change in the abundance of burst-capable neurons does not represent a shift in the underlying rhythmogenic elements of the network. Indeed, blockade of synaptic inhibition in the preBötC changes breathing pattern but does not eliminate rhythmogenesis *in vitro* or *in vivo* (Baertsch et al., 2018; Janczewski et al., 2013), which is inconsistent with a developmental shift towards a distinct, reciprocal inhibition-based-rhythmogenic mechanism. Instead, our study predicts that rhythmogenic elements are conserved, but an increasing amount of excitatory drive becomes required for rhythmogenesis as neurodevelopment progresses (Fig. 6), which may contribute to the difficulties associated with generating rhythmic preBötC slices beyond this early developmental period.

Collectively, these factors may help explain the lack of evidence for intrinsically bursting preBötC neurons *in vivo*. However, a major conclusion of our study is that intrinsic bursting is not a prerequisite for *I*_*NaP*_-dependent rhythmogenesis (Figs. 2 & 4). Therefore, even if preBötC neurons are not capable of intrinsic bursting *in vivo*, this does not indicate that *I*_*NaP*_ isn’t an essential feature of preBötC rhythmogenesis. To the contrary, the conceptual insights of our study support the hypothesis that the preBötC utilizes the same cellular and network features for rhythm generation *in vivo* vs. *in vitro*, during hypoxia, and at different stages of neurodevelopment. However, the network is able to developmentally and/or conditionally alter the abundance of intrinsic bursting/pre-inspiratory spiking phenotypes, characteristics of the network rhythm (frequency, amplitude, shape), and the amount of excitability required for rhythmogenesis. Conditions associated with less intrinsic bursting and more pre-inspiratory spiking generally result in a more dynamic preBötC network that is able to produce a wider range of frequencies with relatively small changes in excitatory input (Figs. 2,6,7). From a teleological perspective, it may make sense for activity phenotypes of preBötC neurons to transition away from intrinsic bursting as development progresses and breathing becomes integrated with an increasingly complex repertoire of non-respiratory behaviors such as sniffing, vocalizing, nociception, emotion, and swallowing (Arthurs et al., 2023; Phillips et al., 2012; Liu et al., 2021; Chiang et al., 2019; Huff et al., 2023). Breathing *in vivo* must also be easily stopped and started, e.g. breath hold, and such changes in the preBötC may ensure it continues to operate near this phase transition with sufficient gain to allow optimal responses to internal and external inputs, consistent with *the critical brain hypothesis* (Hesse and Gross, 2014).

The parameters of our model are based on available data. However, computational models are always limited by approximations and cannot include all biological variables. For example, we do not know how each individual conductance scales with development, or whether time constants for each voltage-gated parameter are similarly affected by temperature. However, the important conceptual takeaways hold true across different permutations and simulations. 1) Due to interaction with the voltage-dependent properties of *I*_*NaP*_, *anything* that alters spike shape can influence intrinsic bursting. 2) Interacting cellular (*g*_*NaP*_ and excitability) and network (excitatory synaptic interactions) properties form the inexorable substrate for rhythm generation, whereas the activity patterns of individual neurons are conditional phenotypes that reflect changes in network “states” rather than changes in rhythmogenic mechanism. And 3) Artificial and/or physiological effects on spike shape can have important consequences for rhythm characteristics and network flexibility. Because *I*_*NaP*_ is widely expressed in the brain (Su et al., 2001; Brumberg et al., 2000; Alzheimer et al., 1993; Taddese and Bean, 2002) and is a feature of many CPGs (Brumberg et al., 2000; Alzheimer et al., 1993; Taddese and Bean, 2002), the impact of these findings is not limited to respiratory rhythm generation. For example, in locomotor circuits, *I*_*NaP*_-dependent intrinsic bursting is thought to contribute to rhythm generation (Tazerart et al., 2008), and blocking the M-current reduces the spike AHP and converts tonic neurons into *I*_*NaP*_-dependent intrinsic bursters (Verneuil et al., 2020). In the basal ganglia, elevated [*K*^+^]_*ext*_ or loss of dopaminergic inputs decreases spike AHP, which coincides with the emergence of intrinsic bursting and pathological network oscillations (Strauss et al., 2008). In cortical neurons (Brumberg et al., 2000; van Drongelen et al., 2006), *I*_*NaP*_ expression, intrinsic bursting, and network mechanisms are implicated in the generation of oscillations linked to slow-wave sleep, epileptiform activity, and mental disorders such as schizophrenia and autism (Wang, 2010; Sanchez-Vives and McCormick, 2000; Stafstrom, 2007). Thus, the conceptual insights of our study may provide a useful framework for understanding many different forms of brain rhythmicity.

## METHODS AND MATERIALS

### Neuron Model

Model preBötC neurons include a single compartment and incorporate Hodgkin-Huxley style conductances adapted from previously described models (Jasinski et al., 2013; Phillips et al., 2019; Phillips and Rubin, 2019) and/or experimental data as detailed below. The membrane potential of each neuron is governed by the following differential equation:

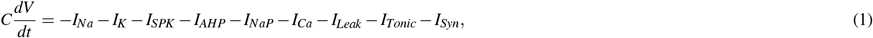

where *C* = 36 *pF* is the membrane capacitance and each *I*_*i*_ represents a current, with *i* denoting the current’s type. The currents include the action potential generating Na^+^ and delayed rectifying K^+^ currents (*I*_*Na*_ and *I*_*K*_), a high voltage activated Na^+^ and K_+_ currents for augmenting spike height (*I*_*SPK*_) and AHP (*I*_*AHP*_), a persistent Na^+^ current (*I*_*NaP*_), voltage-gated Ca^2+^ current (*I*_*Ca*_), K^+^ dominated leak current (*I*_*Leak*_), a tonic excitatory synaptic current (*I*_*Tonic*_) and a dynamic excitatory synaptic current (*I*_*Syn*_) which mediates preBötC network interactions. The currents are defined as follows:

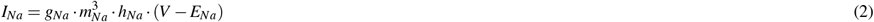

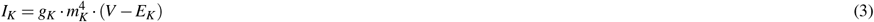

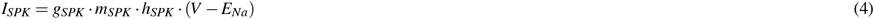

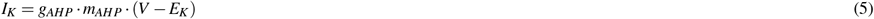

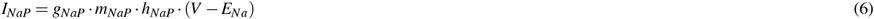

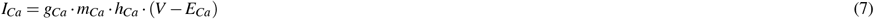

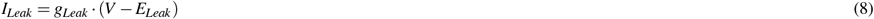

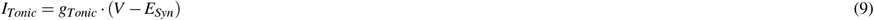

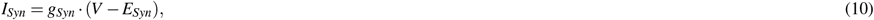

where *g*_*i*_ is the maximum conductance, *E*_*i*_ is the reversal potential, and *m*_*i*_ and *h*_*i*_ are gating variables for channel activation and inactivation for each current *I*_*i*_. The glutamatergic synaptic conductance *g*_*Syn*_ is dynamic and is defined below (Eq. 18). The values used for the *g*_*i*_ and *E*_*i*_ appear in Table 1.

**Table 1.**
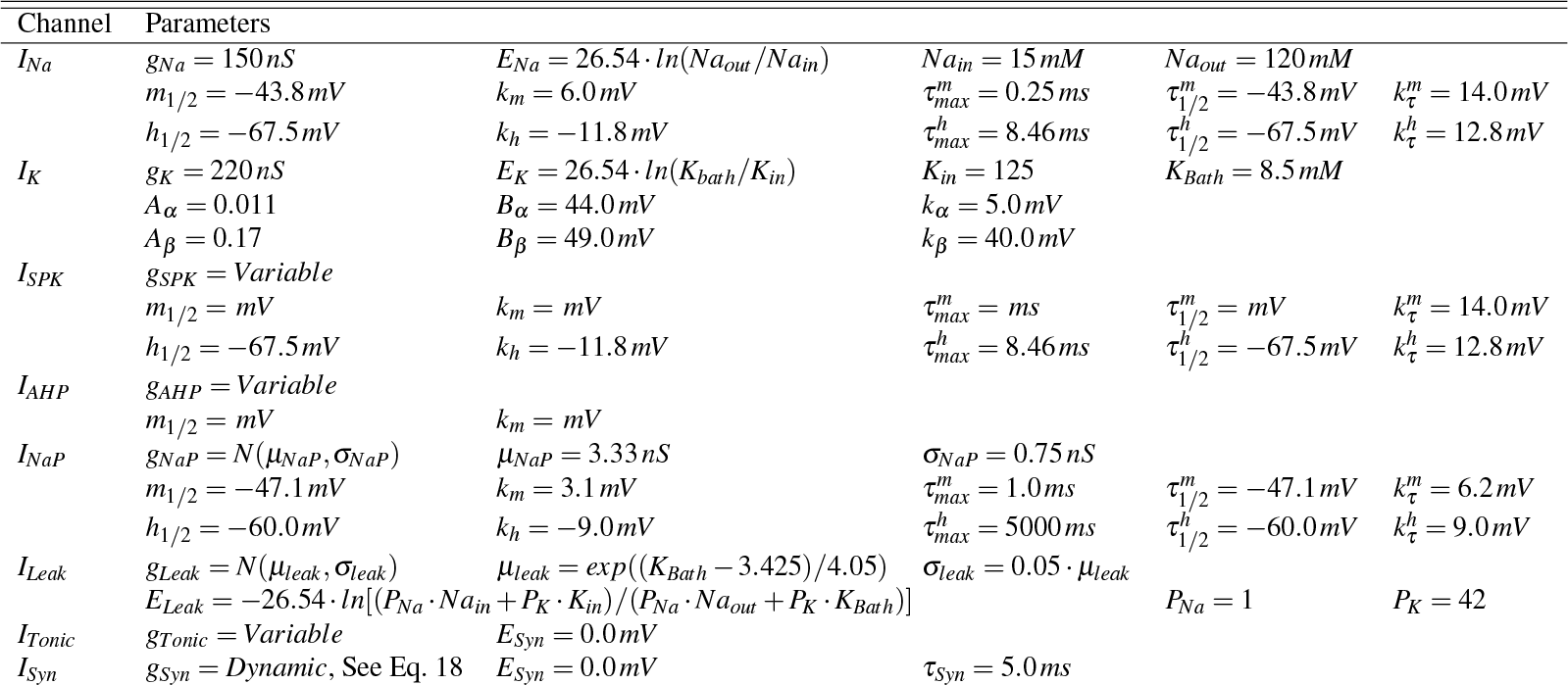
Ionic Channel Parameters.

Activation (*m*_*i*_) and inactivation (*h*_*i*_) of voltage-dependent channels are described by the following differential equation:

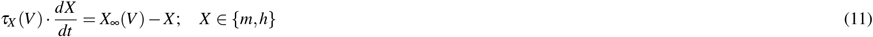

where *X*_∞_ represents steady-state activation/inactivation and *τ*_*X*_ is a time constant. For *I*_*Na*_, *I*_*NaP*_, *I*_*Ca*_, *I*_*SPK*_, and *I*_*AHP*_, the functions *X*_∞_ and *τ*_*X*_ take the forms

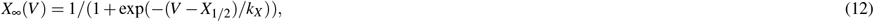

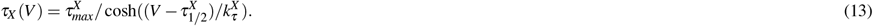

The values of the parameters (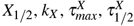, and 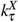) corresponding to *I*_*Na*_, *I*_*NaP*_,*I*_*Ca*_, *I*_*SPK*_ and *I*_*AHP*_ are given in Table 1.

For *I*_*K*_, steady-state activation 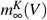 and time constant 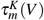 are given by the expressions

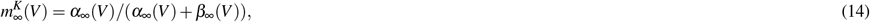

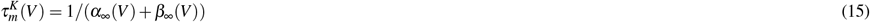

where

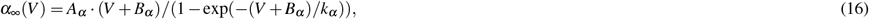

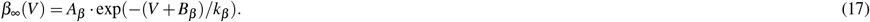

The values for the constants *A*_*α*_, *A*_*β*_, *B*_*α*_, *B*_*β*_, *k*_*α*_, and *k*_*β*_ are also given in Table 1.

When we include multiple neurons in the network, we index them with subscripts. Then the total synaptic conductance (*g*_*Syn*_)_*i*_ of the *i*^*th*^ target neuron is described by the following equation:

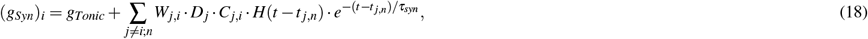

where *W*_*j,i*_ represents the weight of the synaptic connection from neuron *j* to neuron *i, D*_*j*_ is a scaling factor for short-term synaptic depression in the presynaptic neuron *j* (described in more detail below), *C*_*j,i*_ is an element of the connectivity matrix (*C*_*j,i*_ = 1 if neuron *j* makes a synapse with neuron *i* and *C*_*j,i*_ = 0 otherwise), *H*(.) is the Heaviside step function, and *t* denotes time. *τ*_*Syn*_ is an exponential synaptic decay constant, while *t* _*j,n*_ is the time at which the *n*^*th*^ action potential generated by neuron *j* reaches neuron *i*.

This model includes short-term synaptic depression motivated by experimental observations in the preBötC (Kottick and Del Negro, 2015) and past computational models have suggested (Rubin et al., 2009; Guerrier et al., 2015). Synaptic depression in the *j*^*th*^ neuron (*D*_*j*_) was simulated using an established mean-field model of short-term synaptic dynamics (Abbott et al., 1997; Dayan and Abbott, 2001; Morrison et al., 2008) as follows:

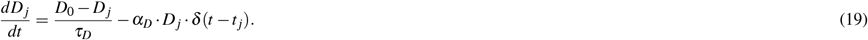

Where the parameter *D*_0_ = 1 sets the maximum value of *D*_*j*_, *τ*_*D*_ = 1000 *ms* sets the rate of recovery from synaptic depression, *α*_*D*_ = 0.2 sets the fractional depression of the synapse each time neuron *j* spikes and *δ* (.) is the Kronecker delta function which equals one at the time of each spike in neuron *j* and zero otherwise. Parameters were chosen to qualitatively match data from Kottick and Del Negro (2015).

### Network construction

The preBötC network was constructed with random synaptic connectivity distribution where the connection probability of *P*_*Syn*_ = 13% as motivated by available experimental estimates Rekling et al. (2000). The weights of excitatory conductances were uniformly distributed such that *W*_*j,i*_ = *U* (0,*W*_*Max*_), where *W*_*Max*_ = 0.2 *nS* is the maximal synaptic conductance.

Heterogeneity of intrinsic cellular properties was introduced into the network by normally distributing the parameters *g*_*leak*_ and *g*_*NaP*_ (Table 1) as well as by uniformly distributing *g*_*SPK*_ in Figs. 4-7 to introduce spike height variability. The *leak* and *NaP* conductances were conditionally distributed in order to achieve a bivariate normal distribution, as suggested by Del Negro et al. (2002a); Koizumi and Smith (2008). In our simulations, this was achieved by first normally distributing *g*_*NaP*_ in each neuron according to the values presented in Table 1. Then a property of bivariate normal distribution was used which says that the conditional distribution of *g*_*leak*_ given *g*_*NaP*_ is itself a normal distribution with mean 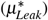 and standard deviation 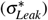 described as follows:

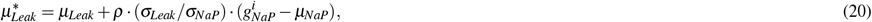

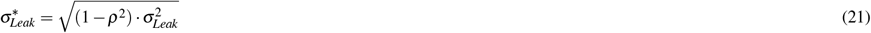

In these equations, *μ*_*Leak*_ and *μ*_*NaP*_ are the mean and *σ*_*Leak*_ and *σ*_*NaP*_ are the standard deviation of the *g*_*Leak*_ and *g*_*NaP*_ distributions, while *ρ* = 0.8 represents the correlation coefficient and 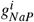 represents the persistent sodium current conductance for the *i*^*th*^ neuron. All parameters are given in Table 1.

### Simulating temperature dependent changes in gating time constants and membrane capacitance

The rate constants for channel gating change exponentially with temperature and is characterized by a Q10 temperature coefficient, which is a measure of the degree to which the rate of a biological process depends on temperature over 10^*°*^C (Sterratt, 2015). Q10 values commonly observed for rate constants of voltage-dependent gating dynamics typically range from 1 to 3 (Matteson and Armstrong, 1982; Collins and Rojas, 1982; Fohlmeister et al., 2010; Yu et al., 2012). For simplicity and feasibility of these experiments, we assumed a Q10 of 1.5 in all voltage-dependent channel rate constants (Yu et al., 2012; Caplan et al., 2014). The resulting scaling factor (Fig. 7 Supplement 1B) was then multiplied by all of the time constants of the voltage-dependent gating variables (*τ*_*X*_ (*V*), Eq. 13) as well as the time constants for the synaptic current (*τ*_*syn*_ in Eq. 18) and the rate of recovery from synaptic depression (*τ*_*D*_, Eq. 19). In addition to changes in rate constants, cells also experience a temperature-dependent increase in surface area, leading to changes in capacitance (Shapiro et al., 2012; Pinto et al., 2021, 2022) at a rate of approximately 0.3% per ^*°*^C (Plaksin et al., 2018). As such, the model membrane capacitance was increased at a rate of 0.3% per ^*°*^C (see Fig. 7 & Fig. 7 Supplement 1C).

### Data analysis and definitions

Data generated from simulations was post-processed in MATLAB software ver. R2020b (MathWorks, Natick, MA, USA). An action potential was defined to have occurred in a neuron when its membrane potential *V*_*m*_ increased through − 35*mV*. Histograms of population activity were calculated as the number of action potentials per 20 *ms* bin per neuron, with units of Hz. The amplitudes and frequency of network rhythms were determined by first identifying the peaks and then calculating the inverse of the interpeak interval from the population histograms. Quantification of spike height and AHP as a function of *g*_*SPK*_, *g*_*AHP*_, or other parameter manipulations (as in Figs. 5–7) was done with *g*_*NaP*_ = 0 *nS* to eliminate intrinsic bursting which would make quantification of AHP impossible. To quantify the percentage of the population that became active since the prior burst we counted the number of neurons in the population that spiked starting 500 *mS* after the peak of one burst to 500 *ms* after the peak of the next burst, except in cases where the burst duration was longer than 500 *ms* in which case this window was manually extended.

### Integration methods

All simulations were performed locally on an eight-core computer running the Ubuntu 20.04 operating system. Simulation software was custom written in C++ and compiled with g++ version 9.3.0. Numerical integration was performed using the first-order Euler method with a fixed step-size (∆*t*) of 0.025*ms*. All model codes will be made freely available GitHub upon publication of this work.

## DATA AND CODE AVAILABILITY

Original code will be posted on GitHub and publicly available upon publication of this manuscript.

## ACKNOWLEDGMENTS AND FUNDING SOURCES

This work was supported by National Institutes of Health grants R01 HL166317 (N.A.B), R00 HL145004 (N.A.B) and K01 1K01DA058543-01 (R.S.P).

## SUPPLEMENTARY MATERIAL

**Figure 1 Supplement 1.**
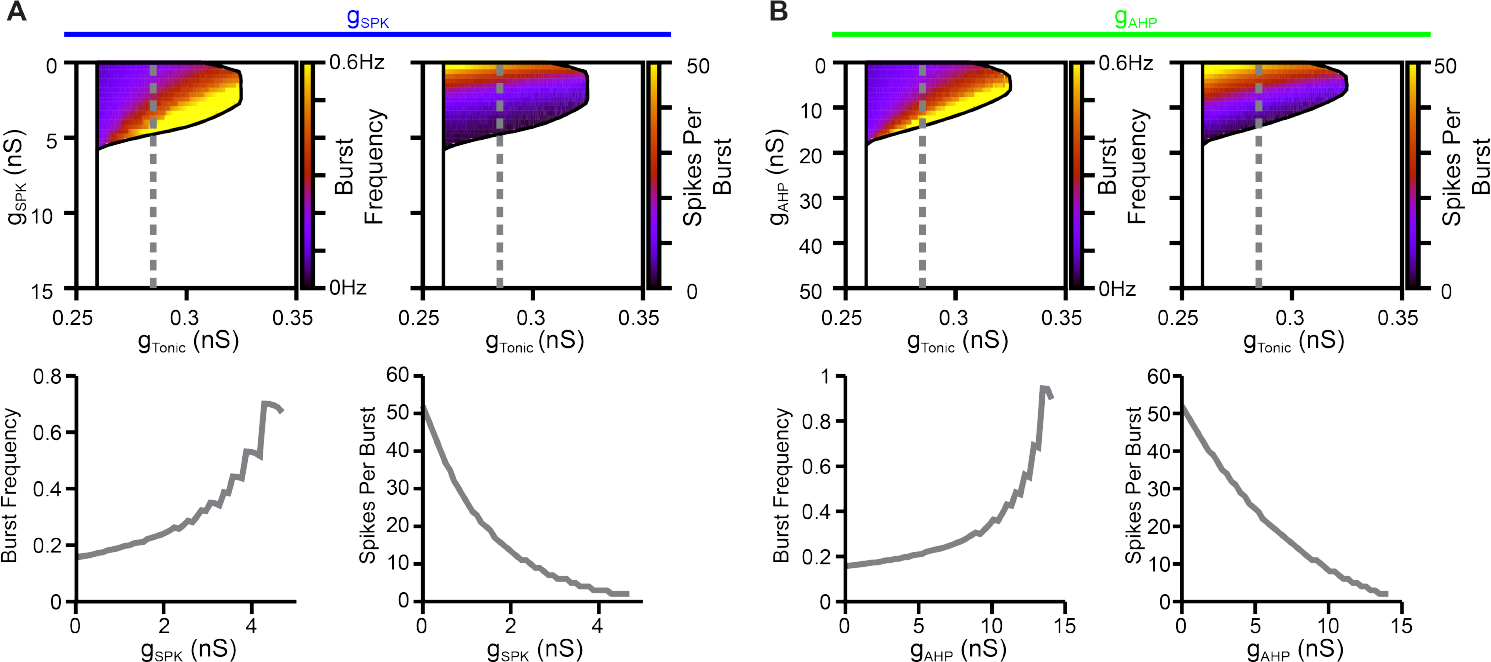
Effect of changes in (A) *g*_*SPK*_ or (B) *g*_*AHP*_ on burst frequency (left) and the number of spikes per burst (right).

**Figure 2-Figure Supplement 1.**
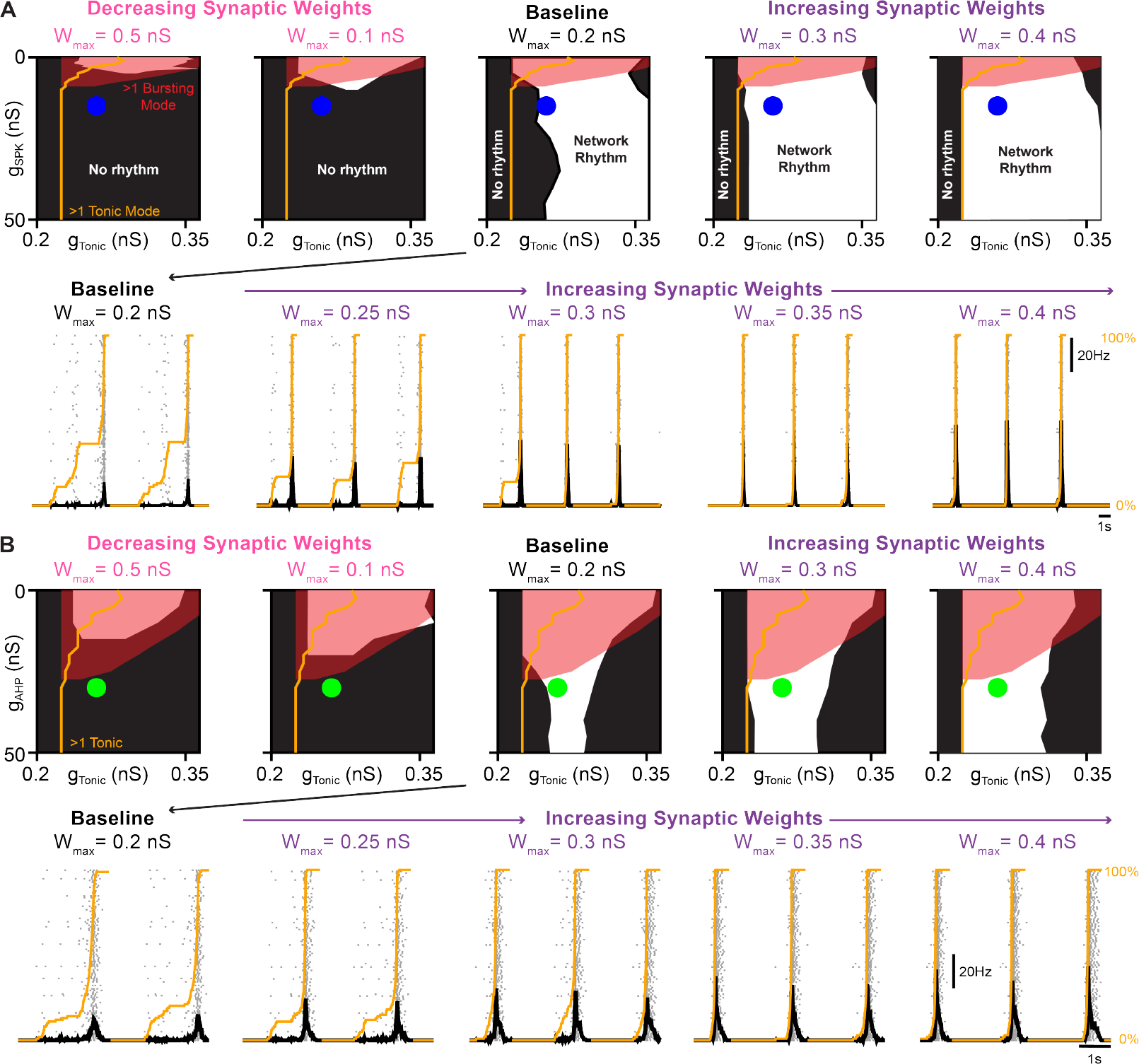
Interactions between spike shape, intrinsic bursting, and synaptic weight for network rhythmogenesis. In networks with (A) altered *g*_*SPK*_ or (B) altered *g*_*AHP*_ the parameter space supporting network rhythmogensis (white regions) was collapsed by decreasing synaptic weights and expanded by increasing synaptic weights. Blue (*g*_*AHP*_ = 30 *nS*) and green (*g*_*SPK*_ = 15 *nS*) dots correspond to *g*_*SPK*_/*g*_*AHP*_ and *g*_*Tonic*_ values of representative traces at baseline and during increasing synaptic weight. Orange lines in example traces indicate the percentage of neurons in the network that have become active since the preceding network burst.

**Figure 2-Figure Supplement 2.**
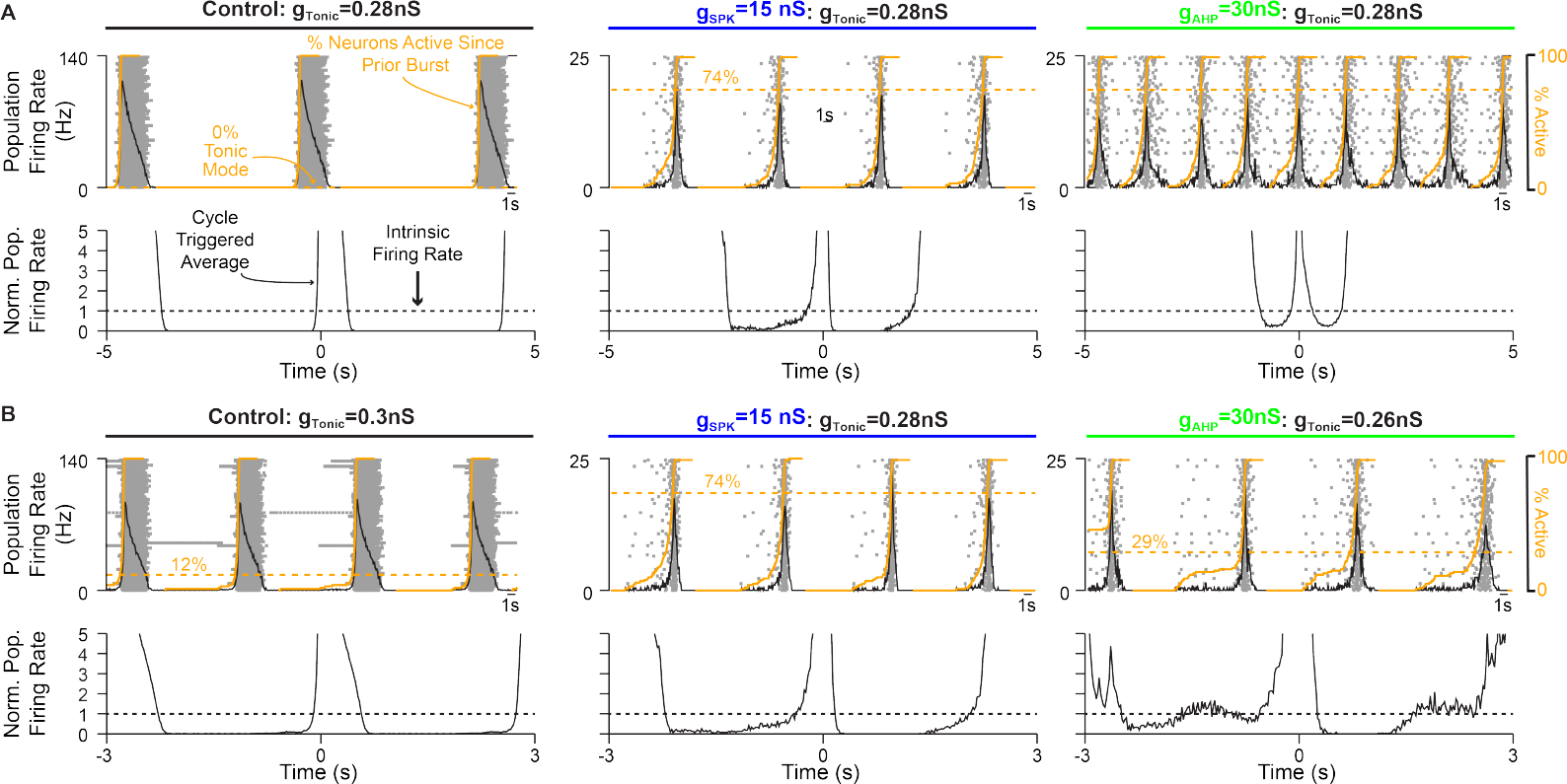
Relationship between pre-inspiratory spiking, the percentage of neurons in tonic spiking mode and the intrinsic network firing rate. Example traces (top) and cycle triggered averages (bottom) in networks with (A) fixed excitability (*g*_*Tonic*_) or (B) altered excitability such that network frequencies are roughly equal (≈ 3 *Hz*). Notice the emergence of pre-inspiratory spiking coincides with the transition of neurons into tonic mode due in the control network and in networks with altered spike shapes.

**Figure 3 Supplement 1.**
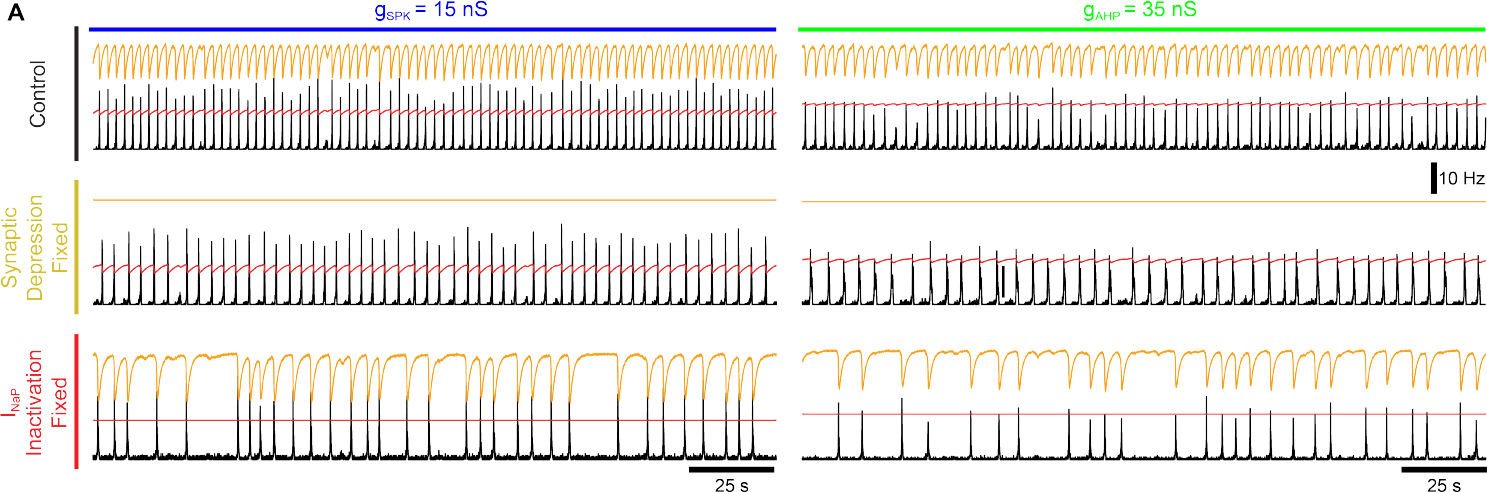
Example network activity (firing rate) and corresponding synaptic depression (orange lines) and *I*_*NaP*_ inactivation (red lines) in networks with *g*_*SPK*_ = 15 *nS* (left) or *g*_*AHP*_ = 35 *nS* (right) under baseline conditions (top) or after fixing synaptic depression (middle) or *I*_*NaP*_ inactivation (bottom).

**Figure 4 Supplement 1.**
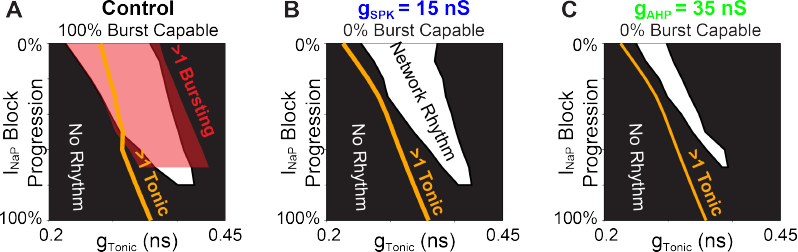
Parameter space supporting intrinsic bursting (red) and network rhythmogenesis (white) as a function of excitability (*g*_*Tonic*_) during progressive *I*_*NaP*_ block in (A) a control network with 100% of neurons initially burst capable (*g*_*SPK*_ = *g*_*AHP*_ = 0) and in networks with (B) *g*_*SPK*_ = 15 *nS* or (C) *g*_*AHP*_ = 35*nS* to eliminate intrinsic bursting. Orange lines indicate *g*_*Tonic*_ value at which ≥ 1 neuron enters tonic spiking mode.

**Figure 4 Supplement 2.**
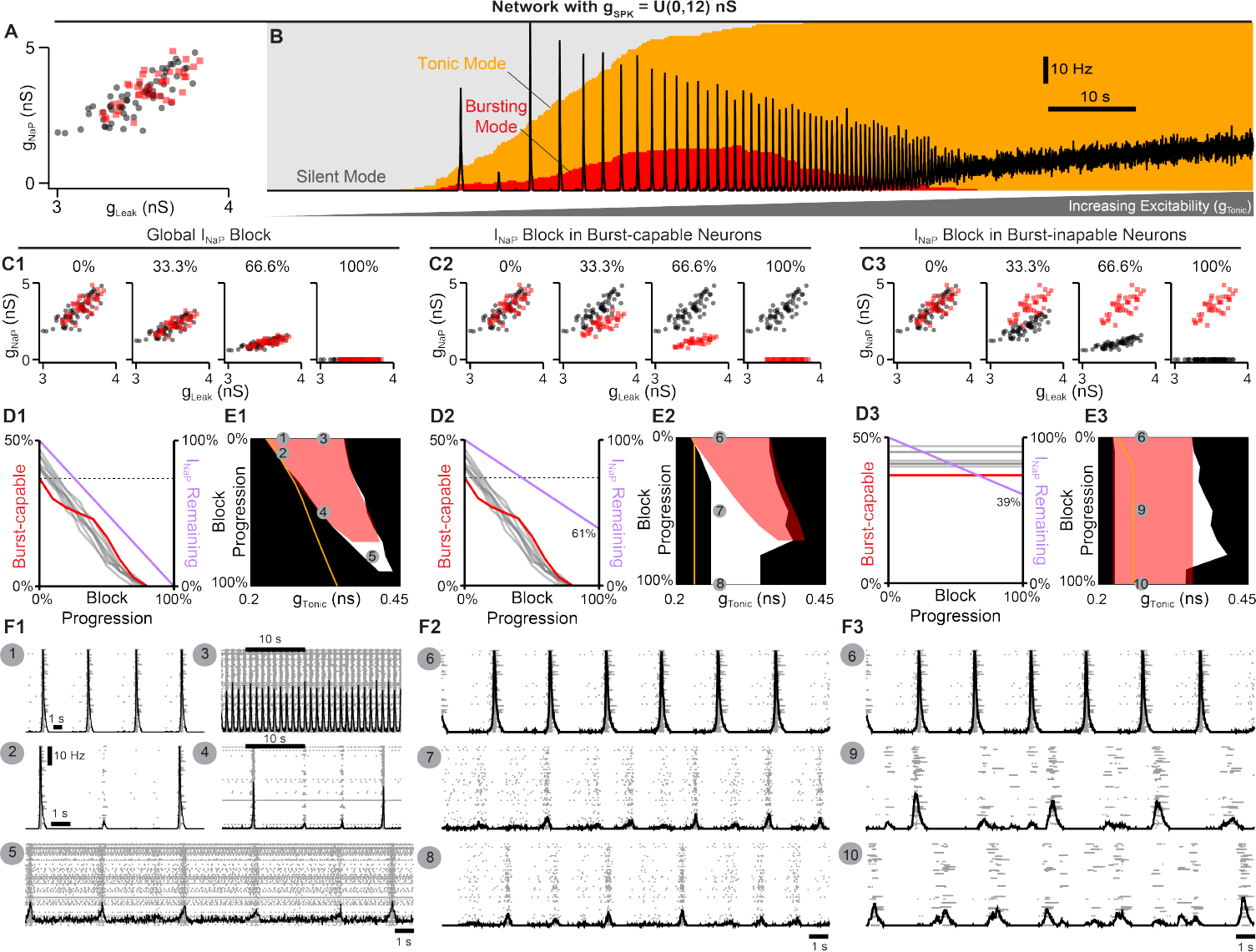
Selective block of *I*_*NaP*_ in burst-capable or burst-incapable neurons has similar consequences for rhythm generation. (A) Distributions of *g*_*NaP*_ and *g*_*Leak*_ among burst-capable (red) and incapable (black) neurons in a network with *g*_*SPK*_ = *U* (0, 12) *nS*. (B) Prevalence of silent, bursting, and tonic intrinsic cellular activities with overlaid network firing rate during increasing *g*_*Tonic*_ in the same network. (C1-3) Comparison of global *I*_*NaP*_ block (C1) vs. progressive *I*_*NaP*_ block specifically in neurons that are initially burst-capable (C2) or burst-incapable (C3). (D1-3) Fraction of the network that is burst-capable and amount of *I*_*NaP*_ remaining as a function of *I*_*NaP*_ block progression. (E1-3) Parameter space supporting intrinsic bursting (red) and network rhythmogenesis (white) as a function of excitability (*g*_*Tonic*_) during progressive *I*_*NaP*_ block. (F1-F3) Raster plots and overlaid network firing rate corresponding to points 1-10 shown in E1-3.

**Figure 5 Supplement 1.**
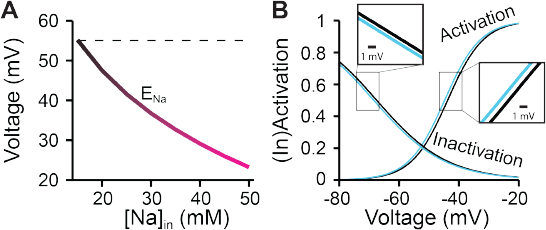
Hypoxia related effects of (A) accumulating [*Na*^+^]_*in*_ on sodium reversal potential and (B) a hyperpolarizing shift in the (in)activation dynamics of spike generating sodium currents.

**Figure 6 Supplement 1.**
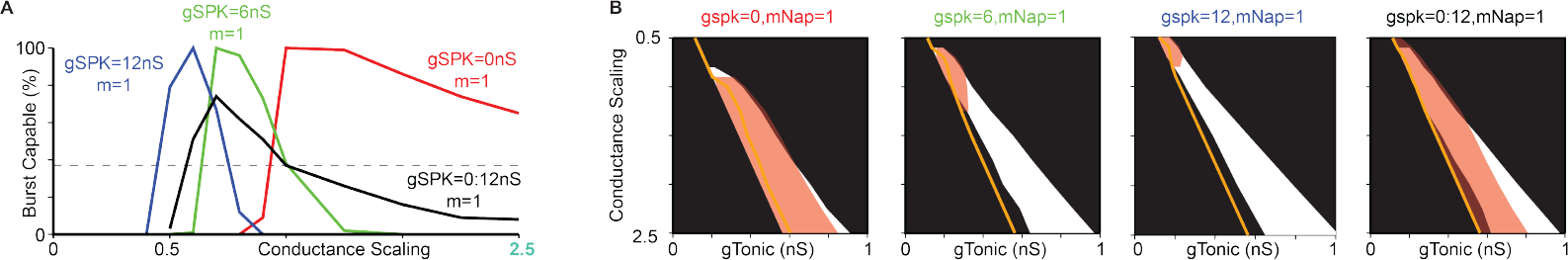
Comparison of conductance scaling across networks with *g*_*SPK*_ = 0 *nS, g*_*SPK*_ = 6 *nS, g*_*SPK*_ = 12 *nS*, or *g*_*SPK*_ = *U* (0, 12) *nS* showing (A) fraction of the network that is burst-capable, and (B) parameter spaces supporting intrinsic bursting (red) and network rhythmogenesis (white) as conductances are up- or down-scaled (Orange lines indicate *g*_*Tonic*_ where ≥ 1 neuron enters tonic spiking mode).

**Figure 6 Supplement 2.**
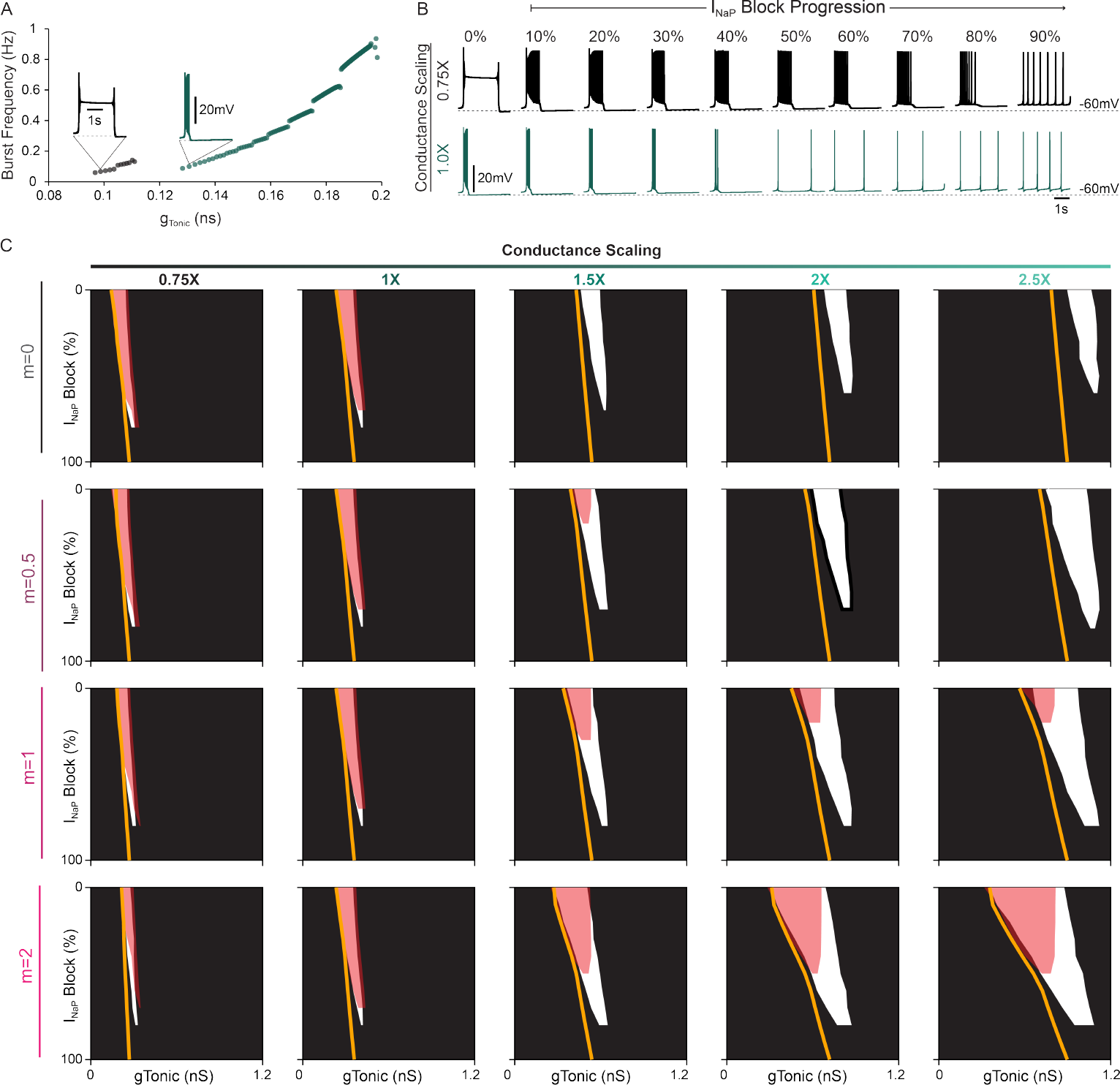
(A) Relationship between excitability (*g*_*Tonic*_) and burst frequency and (B) effect of simulated *I*_*NaP*_ block on intrinsic bursting capabilities for a neuron in with reduced conductance scaling (0.75X,m=1) compared to control scaling (1.0X,m=1). (C) Parameter space supporting network rhythmogenesis during progressive *I*_*NaP*_ block with scaled conductances.

**Figure 7 Supplement 1.**
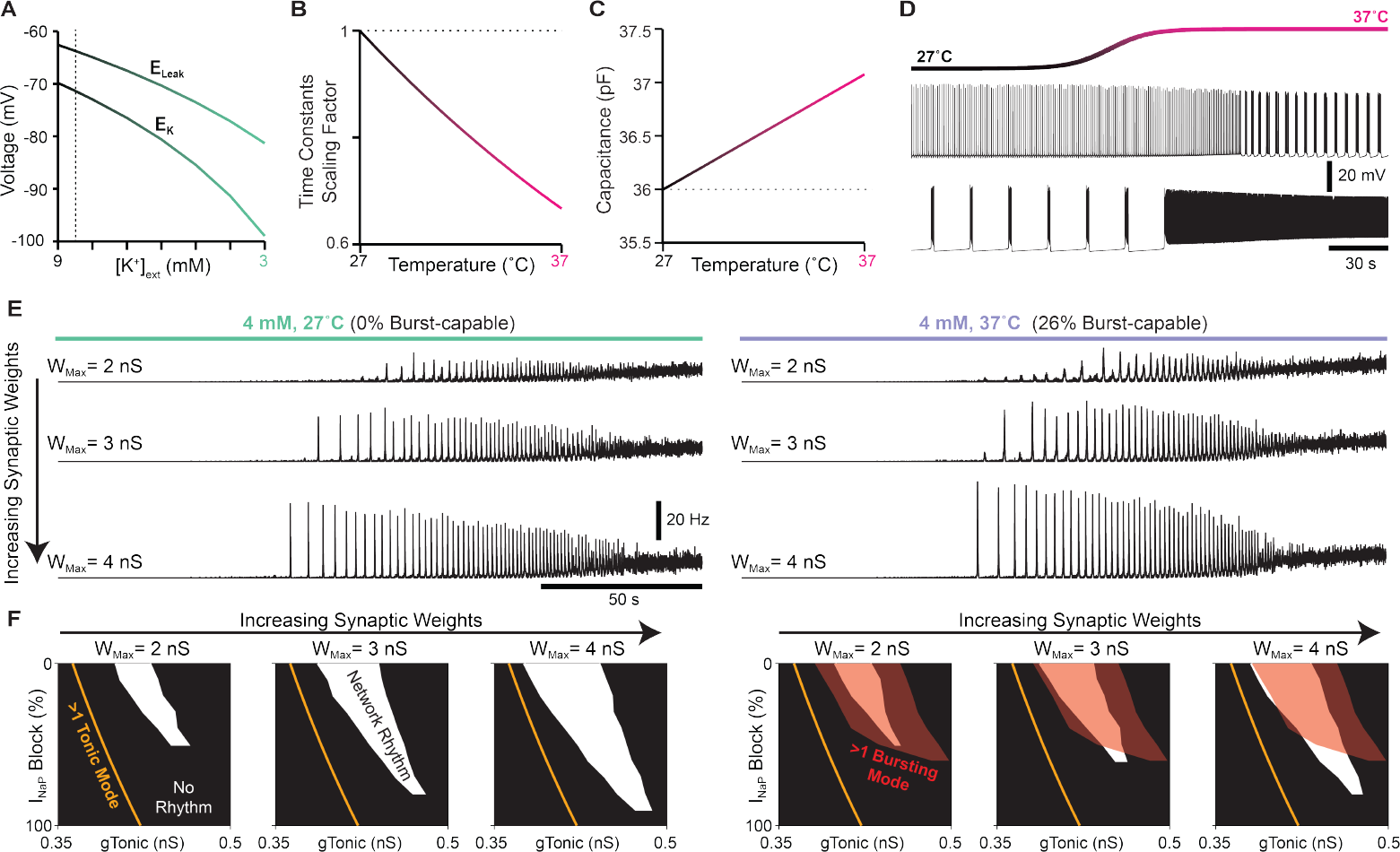
Impact of extracellular potassium, temperature and synaptic weights on network properties and dynamics. (A) Relationship between the potassium (*E*_*K*_) and leak (*E*_*Leak*_) reversal potentials and extracellular potassium [*K*^+^]_*ext*_. Relationship between the scaling of time constants (B) and cellular capacitance (C) and the imposed temperature. (D) Example voltage traces illustrating the transition of a neuron from tonic to bursting mode and from bursting to tonic mode in response to an increase in temperature. (E) Effect of increases in synaptic weights on the network rhythm at physiological potassium and *in vitro* (left) or *in vivo* (right) temperatures. (F) Simulated *I*_*NaP*_ attenuation on network rhythms and intrinsic bursting.

**Figure 7 Supplement 2.**
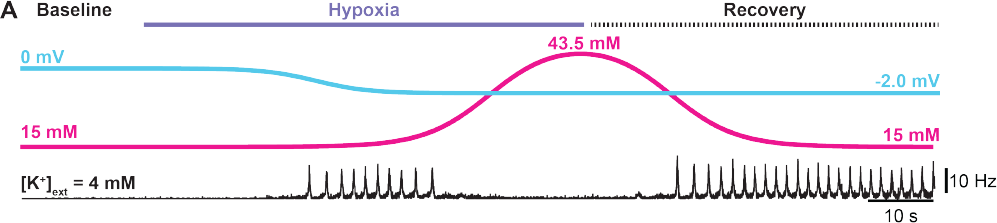
Simulated hypoxia at physiological. [*K*^+^]_*ext*_. (A) Network rhythm during transient hypoxia and recovery.

